# Genome-wide association study and polygenic risk score analysis for hearing measures in children

**DOI:** 10.1101/2020.07.22.215376

**Authors:** Judith Schmitz, Filippo Abbondanza, Silvia Paracchini

## Abstract

An efficient auditory system contributes to cognitive and psychosocial development. A right ear advantage in hearing thresholds (HT) has been described in adults and atypical patterns of left/right hearing threshold asymmetry (HTA) have been described for psychiatric and neurodevelopmental conditions. Previous genome-wide association studies (GWAS) on HT have mainly been conducted in elderly participants whose hearing is more likely to be affected by external environmental factors. We analyzed HT and HTA in a children population cohort (ALSPAC, *n* = 6,743). Better hearing was associated with better cognitive performance and higher socioeconomic status (SES). At the group level, HTA suggested a left ear advantage (mean = -0.28 dB) that was mainly driven by females. SNP heritability for HT and HTA was 0.17 and 0.01, respectively (*n* = 5,344). Genetic correlation analysis confirmed associations between HT, reading ability, listening comprehension, and GCSE scores. GWAS for HT did not yield significant hits but polygenic risk score (PRS) analysis showed significant associations of higher educational attainment (EA, ß = -1564.72, p = .008) and risk for schizophrenia (ß = -241.14, p = .004) with lower HT, i.e. better hearing. In summary, we report new data supporting associations between hearing measures and cognitive abilities at the behavioral level. Genetic analysis suggests shared biological pathways between cognitive and sensory systems and provides evidence for a positive outcome of genetic risk for schizophrenia.

## Introduction

The auditory system involves a series of complex distributed cerebral networks and its impairment affects psychosocial, emotional and cognitive development (Stevenson et al., 2018). Hearing-impaired children are at increased risk for learning disabilities (S. M. R. Choi, Kei, & Wilson, 2020) and even within the normal range, better hearing has been associated with better reading skills, working memory and nonverbal IQ in a sample of 1,638 UK school children (Moore, Zobay, & Ferguson, 2019).

Hearing ability is usually defined as the threshold in decibel (dB) at which a tone is perceived, so that lower values indicate better hearing. An age-related hearing decline is well documented. In a Korean general population sample (*n* > 15,000) the hearing threshold (HT) for medium frequencies declined from 3 dB in adolescents to 38 dB in elderly participants (Park, Shin, Byun, & Kim, 2016). A sex difference in favor of women was found in adults (*n* = 10,145; 30-69 years old), but not in children and young adults (*n* = 3,458).

Successful hearing requires transformation of changes in air pressure into vibrations in the basilar membrane that is transferred onto sensory hair cells of the inner ear, whose depolarization is initiated by deflection of mechano-sensitive hair bundles (Schwander, Kachar, & Müller, 2010). The auditory nerve transmits these signals to the cochlear nucleus in the brainstem. The majority of the input is transmitted to the contralateral superior olivary complex, while a minor part of the input is transmitted ipsilaterally (Felix, Gourévitch, & Portfors, 2018; Warren & Liberman, 1989). Greater contralateral medial olivonuclear suppression in the right compared to the left ear (Khalfa & Collet, 1996) has been suggested as the underlying correlate of a fundamental functional asymmetry between the left and right ear. This hearing threshold asymmetry (HTA) has typically been reported as HT left – HT right so that positive values indicate an advantage of the right and negative values indicate an advantage of the left ear. In a study of more than 50,000 adults Chung, Mason, Gannon, and Willson (1983), a right ear advantage (HTA between 1 dB and 4 dB) has been reported with more pronounced HTA in males than in females. In a children sample of *n* = 1,191, a right ear advantage has been reported, albeit to a smaller extent than in adults (Eagles, 1973). Other authors found a general right ear advantage in males (*n* = ∼400, HTA between 0.1 dB and 0.5 dB) and a left ear advantage in females for specific frequencies (*n* = ∼400, HTA between -0.1 and -0.4 dB) (Kannan & Lipscomb, 1974). Smaller studies reported a general left ear advantage in children (Rahko & Karma, 1989; Roche, Siervogel, Himes, & Johnson, 1978).

An absence of HTA has been reported in schizophrenia (Gruzelier & Hammond, 1979; Mathew, Gruzelier, & Liddle, 1993) and ADHD (Combs, 2002). Moreover, symmetrical contralateral suppression in the olivary complex in the left and the right ear in schizophrenia (Veuillet et al., 2001) is in contrast with the right ear advantage typically found in controls. In children and adolescents, a right ear advantage has been reported in a sample of *n* = 22 with autism spectrum disorder (ASD) while no asymmetry was found in the control group (Khalfa et al., 2001). A developmental effect towards stronger HTA has been reported in controls that was absent in ASD children (*n* = 24) (James & Barry, 1983). Reduced laterality in processing auditory stimuli was reported in ASD and bipolar disorder (BIP) suggesting that HTA is linked to neurodevelopmental disorders (Reite et al., 2009; Schmidt, Rey, Oram Cardy, & Roberts, 2009).

Twin and family studies estimated the heritability for HT to range from .26 to .75 with larger environmental effects for the non-dominant ear (Gates, Couropmitree, & Myers, 1999; Viljanen, Era, et al., 2007; Viljanen, Kaprio, et al., 2007). Genome-wide association studies (GWAS) focused on age-related hearing loss in subjects ranging from 45 to 75 years of age and identified genes including *GRM7* and *ESRRG* (Fransen et al., 2015; Friedman et al., 2009; Huyghe et al., 2008; Nolan et al., 2013). GWAS for normal hearing included subjects from 18 to 92 years of age (Girotto et al., 2011; Vuckovic et al., 2015; Wolber et al., 2014) and implicated several genes (i.e. *DCLK1, PTPRD, GRM8, CMIP, SIK3, PCDH20, SLC28A3*), some of which have been associated with neurodevelopmental traits (Håvik et al., 2012; Newbury et al., 2009; Newbury et al., 2019; Scerri et al., 2011; Takaki et al., 2004; Zhang et al., 2014). A case control design based on electronical health records on age-related hearing loss identified SNPs near *ISG20* and *TRIOBP* (Hoffmann et al., 2016), which had previously been associated with prelingual nonsyndromic hearing loss (Diaz-Horta et al., 2012). In the UK Biobank, 41 and 7 independent loci have been identified for hearing difficulty and hearing aid use, respectively, implicating genes such as *CDH23, EYA4, KLHDC7B* and again *TRIOBP* (Wells et al., 2019). Mutations in *CDH23* have been associated with early-onset hearing loss and Usher syndrome causing early-onset deafness (Schultz et al., 2011).

However, older subjects have had more exposure to environmental factors which might affect hearing, such as extensive noise (Stanbury, Rafferty, & Rosenman, 2008), medication (Ruhl, Cable, & Martell, 2014), chemicals (Morata, 2003) and medical conditions (Kakarlapudi, Sawyer, & Staecker, 2003). An investigation of HT in 250 monozygotic (MZ) and 307 dizygotic (DZ) twin pairs from 36 to 80 years of age suggests that environmental effects become more significant with age (Karlsson, Harris, & Svartengren, 1997). Despite this age effect, no study has ever investigated genetic factors involved in hearing function in children.

Here, we analyzed hearing measures in children from the Avon Longitudinal Study of Parents and Children (ALSPAC) (*n* = 6,743). Consistent with previous studies we found that better hearing is associated with enhanced cognitive skills and higher SES. We report the first GWAS for HT in children (*n* = 5,344). In addition to single marker trait associations, we conducted gene-based and gene set analysis and tested the effects of polygenic risk scores (PRS) for a range of neurodevelopmental disorders, IQ and educational attainment (EA). Our results suggest that PRS for higher EA and schizophrenia risk are associated with better hearing.

## Materials and Methods

### Cohort

ALSPAC is a longitudinal cohort representing the general population living in the Bristol area. Pregnant women resident in the county of Avon, UK, with expected dates of delivery from 1st April 1991 to 31st December 1992 were invited to take part in the study, resulting in 14,062 live births and 13,988 children who were alive at 1 year of age (Boyd et al., 2013; Fraser et al., 2013). From age seven, all children were invited annually for assessments on a wide range of physical, behavioral and neuropsychological traits. Informed written consent was obtained from the parents after receiving a complete description of the study at the time of enrolment into ALSPAC, with the option to withdraw at any time. Ethical approval for the present study was obtained from the ALSPAC Law and Ethics Committee and the Local Research Ethics Committees. The ALSPAC study website contains details of all the data that is available through a fully searchable data dictionary (http://www.bris.ac.uk/alspac/researchers/data-access/data-dictionary/).

### Phenotypes

Audiometry was performed according to British Society of Audiologists standards. Hearing tests were carried out in a room with minimal external noise. Testing was stopped if the background noise level exceeded 35 dBA. The air conduction threshold, i.e. the lowest intensity in decibels at which a tone is perceived 50% of the time (dBHL, decibel hearing level), was tested using either a GSI 61 clinical audiometer or a Kamplex AD12 audiometer. Lower dBHL values indicate better hearing. For each ear, the air conduction threshold level was tested at 500 Hz, 1 kHz, 2 kHz and 4 kHz. For each frequency, stimuli were first presented on the right and then on the left ear. The average threshold across different frequencies was derived for each ear.

After applying exclusion criteria (supplementary methods), a sample of *n* = 6,743 was available for phenotypic analysis (3,344 females, 3,391 males, 8 missing values for sex, mean age = 7.59 years, SD = 0.32 years).

HT was defined as the average air conduction threshold on the better ear. HTA was defined as the absolute difference in air conduction threshold between the left and right ear. Thus, positive values indicate a right ear advantage, while negative values indicate a left ear advantage. Handedness was assessed in terms of writing hand (5,805 right-handers, 787 left-handers, 151 missing values).

Cognitive skills were assessed using tests for reading ability (Rust, Golombok, & Trickey, 1993), communication skills (Bishop, 1998), listening comprehension (Rust, 1996), short term memory (Gathercole, Willis, Baddeley, & Emslie, 1994), total IQ, verbal IQ, performance IQ (Wechsler, Golombok, & Rust, 1991) and EA measured as capped General Certificate of Secondary Education (GCSE) scores (for detailed descriptions, see supplementary methods). Maternal highest educational qualification during pregnancy was used as a proxy for SES (Rashid et al., 2018). Educational qualification was grouped into ‘ CSE and no education’, ‘ vocational’, ‘ O level’, ‘ A level’ and ‘ Degree’.

Children were assigned to neurodevelopmental and control subgroups as defined for the ALSPAC sample previously (Scerri et al., 2011). We specified subgroups for language impairment (LI) (*n* = 155), reading disability (RD) (*n* = 141), ASD (*n* = 35), ADHD (*n* = 21), comorbidity (*n* = 49) and a control sample matched for sex (*n* = 2,071). The strategies for subgroup assignments are reported in the supplementary methods.

### Genotype quality control (QC) and imputation

Genotypes were generated on the Illumina HumanHap550-quad array at the Wellcome Trust Sanger Institute, Cambridge, UK and the Laboratory Corporation of America, Burlington, NC, US. Standard QC was performed as described elsewhere (Brandler et al., 2013). In total, 9,115 subjects and 500,527 SNPs passed QC filtering. Haplotypes were estimated using ShapeIT (v2.r644). QC-filtered autosomal SNPs were imputed using Impute v3 using the HRC 1.1 reference data panel. Poorly imputed SNPs (Info score < 0.8) and SNPs with low minor allele frequency (MAF < 0.05) were excluded from further analysis.

### Statistical analysis

#### SNP heritability

Genome-wide genotype data were available for 5,344 children with phenotypes (2,691 males, 2,653 females, mean age = 7.58 years, SD = 0.31 years). HT and HTA were inverse rank-transformed to achieve a normal distribution. For both phenotypes, SNP heritability (SNP h^2^) was estimated using variance components analysis in BOLT-REML (Loh, Bhatia, et al., 2015) implemented in BOLT-LMM v2.3.4 (Loh, Tucker, et al., 2015). SNP h^2^ was estimated based on directly genotyped SNPs and using sex and the first two ancestry-informative principal components as covariates.

#### Genetic correlation analysis

BOLT-REML applies variance component analysis for multi-trait modelling to estimate genetic correlations among multiple traits measured on the same set of individuals. We used BOLT-REML to estimate genetic correlations (rg) between HT and the eight cognitive skills used for behavioral analysis using sex and the first two ancestry-informative principal components as covariates.

#### GWAS

Association testing was performed using BOLT-LMM v2.3.4 (Loh, Tucker, et al., 2015) under the standard infinitesimal linear mixed model (LMM) framework specifying sex and the first two ancestry-informative principal components as covariates. Overall, 5,305,352 SNPs that were either directly genotyped or imputed and passed QC were tested for association. The genomic inflation factor (λ) was calculated for all SNPs and revealed no evidence of population structure (HT: λ = 1.02, HTA: λ = 1.00).

#### Annotation and gene mapping

We applied FUMA v1.3.6a (Watanabe, Taskesen, van Bochoven, & Posthuma, 2017) on the GWAS summary statistics. Functional consequences of SNPs were obtained by performing ANNOVAR (Wang, Li, & Hakonarson, 2010) using Ensembl genes (build 92). SNPs were mapped to genes based on positional mapping. Intergenic SNPs were annotated to the closest genes upstream and downstream. Input SNPs were mapped to 18,360 protein-coding genes.

#### Replication analysis

For HT, we specifically tested for replication of SNPs associated with quantitative hearing phenotypes in previous studies [*p* < 10^−5^ (Fransen et al., 2015; Girotto et al., 2011), *p* < 10^−6^ (Vuckovic et al., 2015; Wolber et al., 2014)]. We also tested for replication of SNPs showing genome-wide significance in case-control GWAS on age-related hearing loss [*p* < 5 × 10^−8^ (Hoffmann et al., 2016; Wells et al., 2019)]. This procedure resulted in 644 SNPs showing genome-wide statistical significance, which we selected as pre-defined lead SNPs in FUMA. Of the 644 SNPs, FUMA identified 272 independent SNPs (r^2^ =< 0.6, default setting), which were used to determine boundaries of LD blocks that encompassed a total of 1126 SNPs (r^2^>= 0.6, default setting). Among these 1126 SNPs, 411 were available in our study resulting in a Bonferroni-corrected significance level of 0.05/411 = 1.2 × 10^−4^.

#### Gene-based and gene set analyses

FUMA implements MAGMA v1.08 (de Leeuw, Mooij, Heskes, & Posthuma, 2015) to summarize SNP associations at the gene level (gene-based analysis) and associate the set of genes to biological pathways (gene set analysis). Gene-based *p* values were computed using an F-test in a multiple linear principal components regression while accounting for LD between SNPs. Genome-wide significance was defined as *p* = 0.05/18,360 = 2.7 × 10^−6^. For gene set analysis, MAGMA converts gene-based *p* values into *z* values, which reflect the strength of association. MAGMA performs a competitive gene set analysis, comparing the mean association of genes within a gene set with the mean association of genes not in the gene set while correcting for gene size and density. Gene set *p* values were computed for 7,343 gene ontology (GO) terms for biological processes obtained from MsigDB v5.2. The Bonferroni-corrected significance level was set to 0.05/7,343 = 6.8 × 10^−6^.

#### PRS

PRS analyses were carried out using PRSice 2.3.3 (S. W. Choi & O’ Reilly, 2019). PRSice uses GWAS summary statistics as training GWAS to build PRS, which are then tested as predictors for the phenotype of interest in the target sample (ALSPAC). Psychiatric Genomics Consortium summary statistics were downloaded (https://www.med.unc.edu/pgc/data-index/) for schizophrenia (Schizophrenia Working Group of the Psychiatric Genomics Consortium, 2014), ADHD (Neale et al., 2010), ASD and BIP (Psychiatric GWAS Consortium Bipolar Disorder Working Group, 2011) as these are the psychiatric and neurodevelopmental conditions often found to be associated with reduced laterality (Khalfa et al., 2001; Mathew et al., 1993; Reite et al., 2009; Veuillet et al., 2001). Based on associations of HT with cognitive skills, GCSE scores and SES, we downloaded summary statistics for IQ (Savage et al., 2018) from the Complex Trait Genetics lab website (https://ctg.cncr.nl/software/summary_statistics) and EA (Lee et al., 2018) from the Social Science Genetic Association Consortium (https://www.thessgac.org/data).

SNPs were clumped based on LD (r^2^ >= 0.1) within a 250 kb window. PRS were derived as the weighted sum of risk alleles based on odds ratios or beta values from the training GWAS summary statistics. Sex and first two principal components were included as covariates. Results are presented for the optimal training GWAS *p* value threshold (explaining the highest proportion of phenotypic variance in HT) as well as GWAS *p* value thresholds of 0.001, 0.05, 0.1, 0.2, 0.3, 0.4, 0.5, and 1 (all SNPs included). For optimal training GWAS *p* value thresholds and number of SNPs included in the PRS, see Supplementary Table S1. For six training GWAS, the Bonferroni-corrected significance level was set to 0.05/6 = .0083.

Data preparation and visualization was performed using R v.4.0.0. All analysis scripts are available through Open Science Framework (https://osf.io/gewj2/).

## Results

### Phenotypes

We first analyzed the distribution of HT and HTA in the overall sample (*n* = 6,743) (results for the subset used in the GWAS, *n* = 5,344, are shown in Supplementary Figure S1). HT ranged from -8.75 to 20.00. Mean HT was 6.14 (SD = 3.97) (Figure 1A). Females (*n* = 3,344) showed slightly higher HT (mean = 6.30, SD = 4.06) than males (*n* = 3,391, mean = 5.98, SD = 3.88), *t*_(6709.2)_= -3.31, *p* = .001 (Figure 1B).

**Figure 1:**
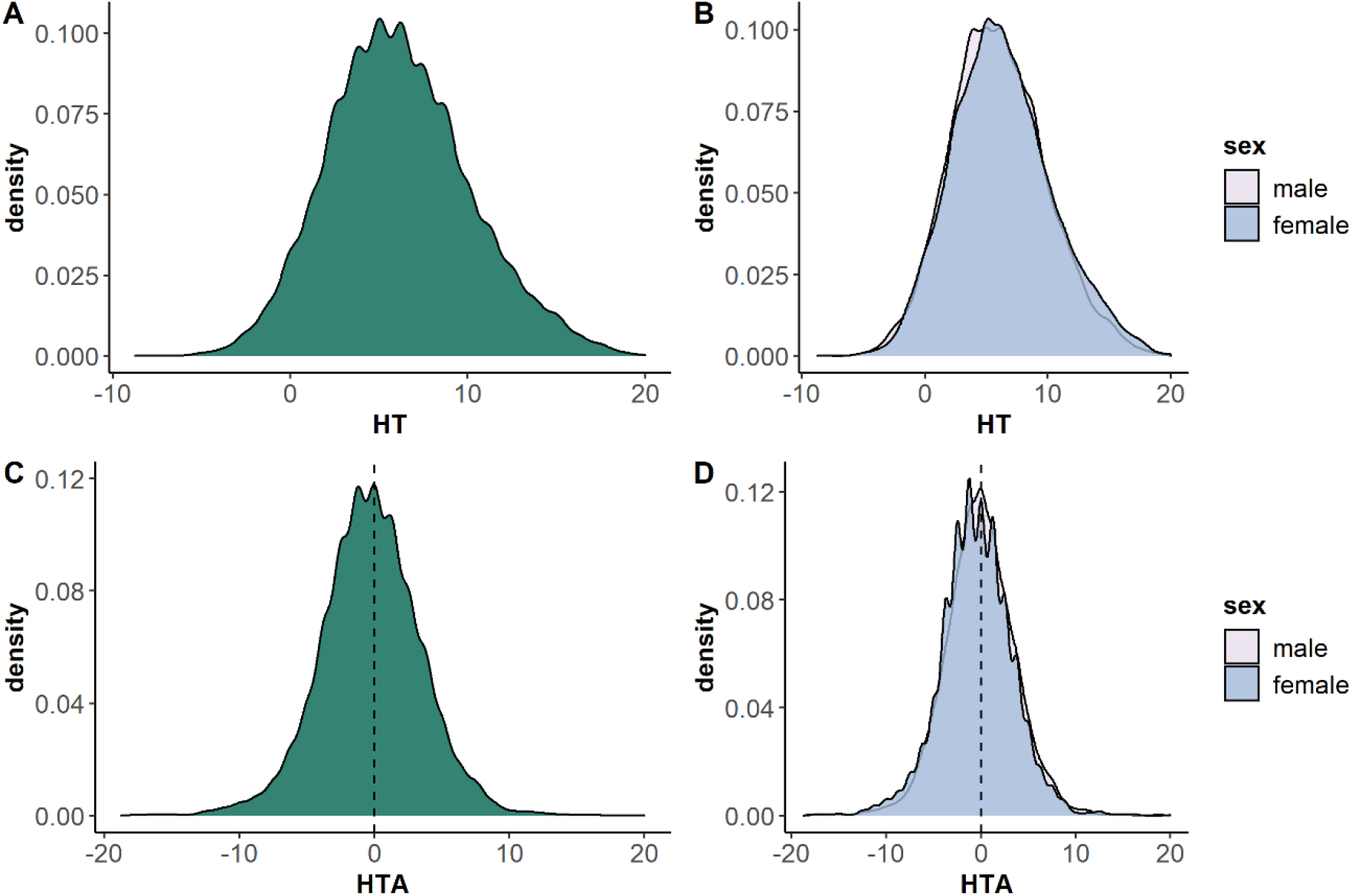
Distribution of HT and HTA. A) Distribution of HT (better ear) in the overall sample (n = 6,743), and B) as a function of sex. C) Distribution of HTA in the overall sample, and D) as a function of sex. The dotted line represents no asymmetry between the left and right ear.

HTA ranged from -18.75 to 20 and revealed a significant left ear advantage for the overall sample (mean = -0.28, SD = 3.75) as determined by one-sample *t*-test against zero (*t*_(6742)_= -6.17, *p* = 7.37 × 10^−10^) (Figure 1C). Females showed stronger HTA (mean = -0.48, SD = 3.81) than males (mean = -0.09, SD = 3.68), *t*_(6716.9)_= 4.33., *p* = 1.54 × 10^−5^ (Figure 1D). We thus performed one-sample *t*-tests against zero for females and males separately. While females showed a significant left ear advantage (*t*_(3343)_ = -7.30, *p* = 3.50 × 10^−13^), there was no ear advantage in males (*t*_(3390)_ = -1.37, *p* = .172). There was no evidence for an effect of handedness on HTA (right-handers: *n* = 5,805, mean = -0.28, SD = 3.75; left-handers: *n* = 787, mean = -0.24, SD = 3.72, *t*_(1015)_ = -0.29, *p* = .776).

Bivariate Pearson correlations revealed significant positive correlations among the different cognitive traits as previously reported (Scerri et al., 2011). There were significant negative correlations after conservative Bonferroni correction (45 comparisons, *p* < .0011) for all cognitive traits but listening comprehension with HT (Figure 2), indicating that lower HT (better hearing) is associated with better cognitive performance (correlation plots are shown in Supplementary Figure S2). There was no association between HTA and cognitive traits.

**Figure 2:**
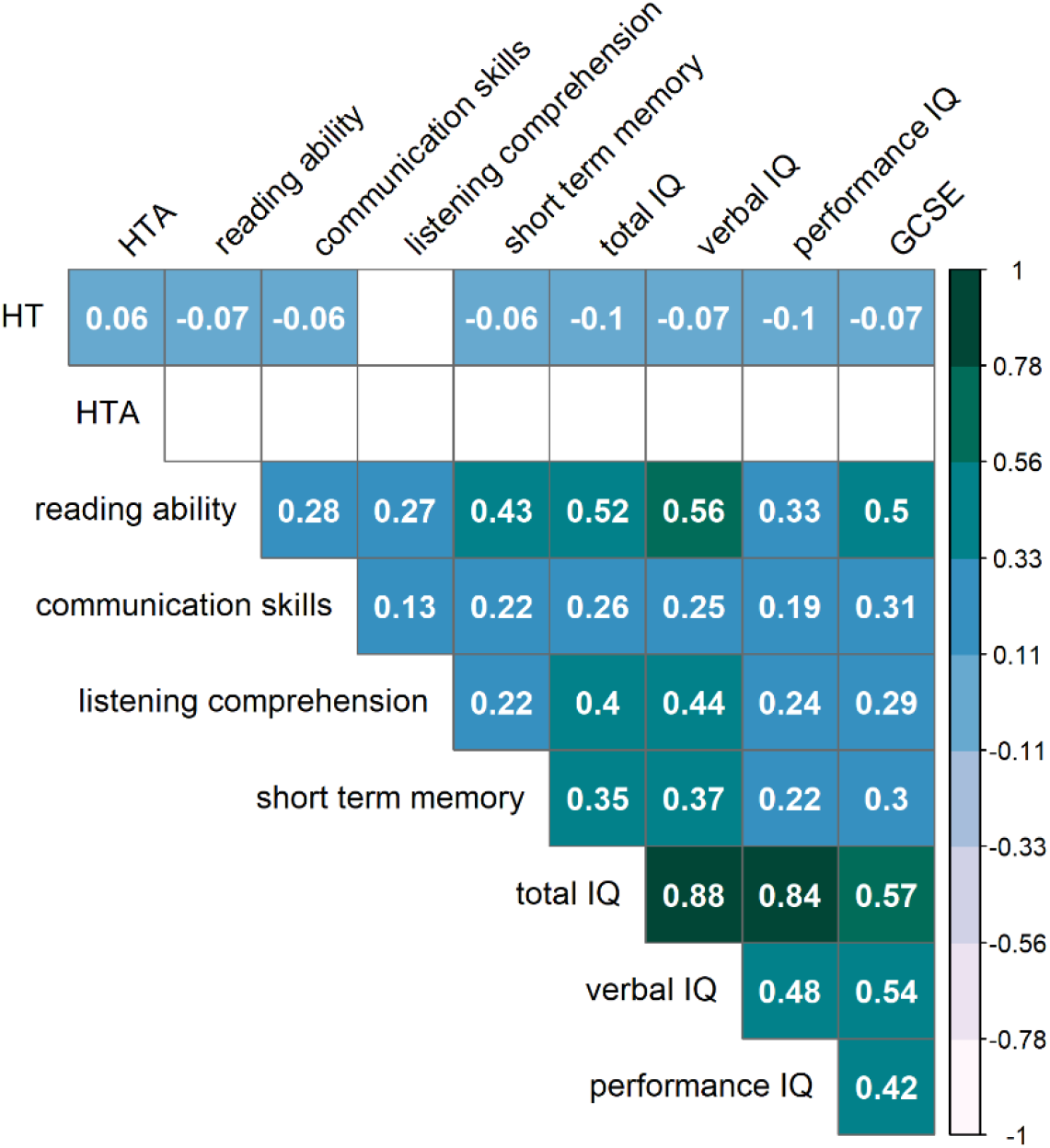
Correlation matrix for HT, HTA and cognitive traits. Correlation coefficients are shown if passing the Bonferroni-corrected significance level (p < .0011). Sample sizes range from n = 4,730 to n = 6,743 depending on data availability.

One-way between-subjects ANOVAs were conducted to compare the effect of SES on HT and HTA. There was a significant effect of SES on HT (*F*_(4,6137)_ = 7.25, *p* = 8.1 × 10^−6^). Post hoc comparisons using the Tukey test indicated significantly lower HT for ‘ O level’ (mean = 6.12, SD = 3.95), ‘ A level’ (mean = 5.91, SD = 3.93) and ‘ Degree’ (mean = 5.92, SD = 4.05) compared to ‘ CSE’ (mean = 6.69, SD = 4.12) and significantly lower HT for ‘ A level’ and ‘ Degree’ compared to ‘ Vocational’ (mean = 6.52, SD = 3.84), indicating higher SES is associated with better hearing (Supplementary Figure S3). There was no significant effect of SES on HTA (*F*_(4,6137)_ = 1.78, *p* = .130).

Two-sample *t*-tests revealed no difference between children affected by neurodevelopmental disorders and sex-matched controls in HT (Supplementary Table S2) or HTA (Supplementary Table S3). However, there was a consistent pattern across neurodevelopmental subgroups with more negative HTA compared to the control group, indicating more leftward asymmetry.

#### SNP heritability

SNP h^2^ was 0.17 (SE = 0.06) for HT and 0.01 (SE = 0.06) for HTA. Because of the extremely low SNP h^2^ for HTA, subsequent genetic analyses were only performed for HT.

#### Genetic correlations

Genetic correlations (rg) between HT and cognitive traits ranged from rg = -.37 (SE = .23) for listening comprehension to rg = .12 (SE = .21) for performance IQ (Figure 3). We observed negative correlations between HT and reading ability, listening comprehension, and GCSE scores, in line with behavioral correlations.

**Figure 3:**
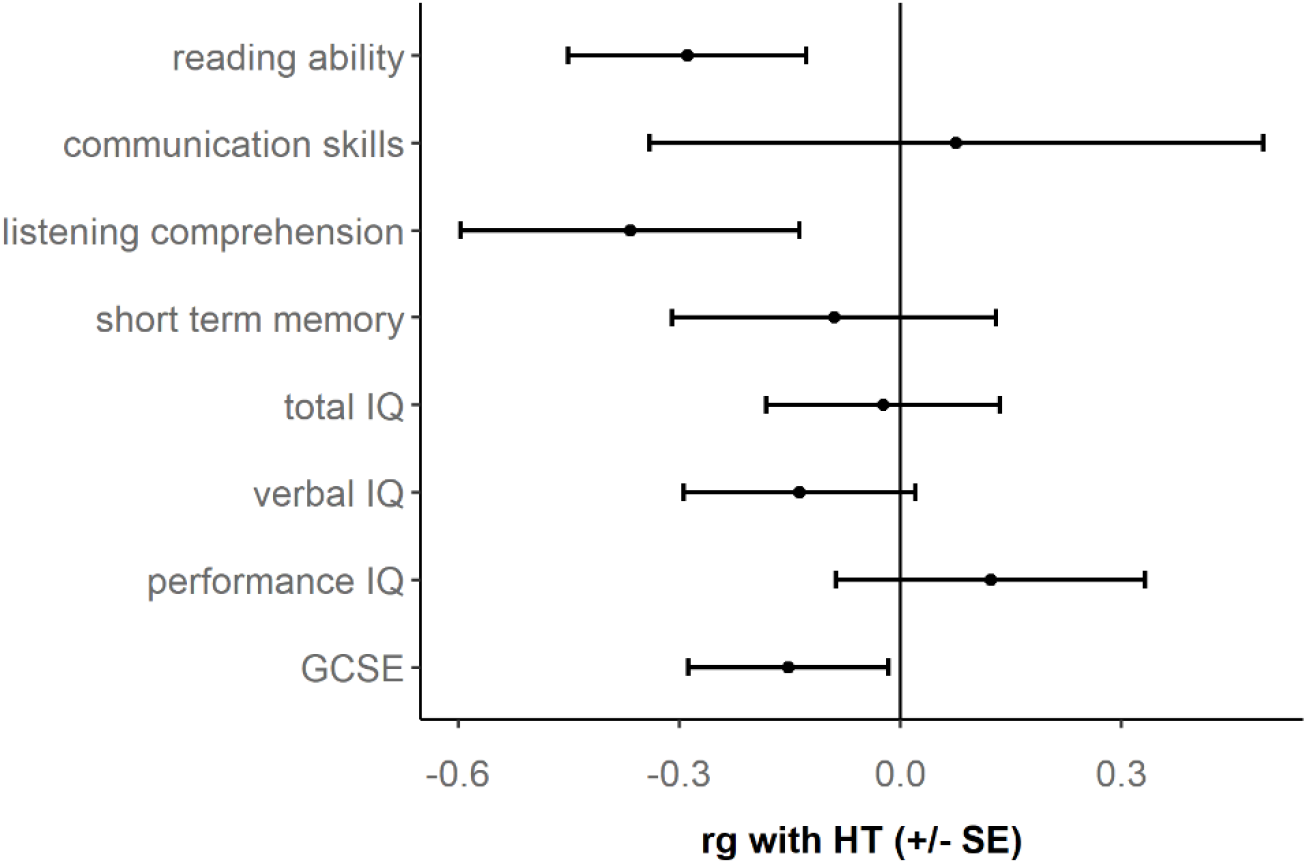
Results of genetic correlation (rg) analysis between HT and cognitive traits.

#### GWAS

GWAS was performed in *n* = 5,344 children. No individual SNP reached genome-wide significance for HT (Supplementary Table S4). The strongest association was found for rs11644235 on chromosome 16 (*p* = 1.8 × 10^−6^) with each copy of the major allele (MAF = 0.42) shifting an individual 0.09 standard deviations towards higher HT (i.e. worse hearing). In gene-based analysis, no gene reached genome-wide significance for HT. Here, the strongest association was found for *AE000662*.*92* (*p* = 5.6 × 10^−5^). Manhattan and QQ plots for SNP-based and gene-based GWAS on HT are shown in Supplementary Figures S4-S7.

In the replication analysis of SNPs derived from previous literature, one SNP reached Bonferroni-corrected significance (rs12955474, ß = 0.17, *p* = 8.1 × 10^−5^, Supplementary Table S5). This SNP had been previously reported in a GWAS for the third PC on HT over different frequencies (0.25, 0.5, 1, 2, 3, 4, 6, 8 kHz) (ß = -0.10, *p* = 3.57 × 10^−7^) (Fransen et al., 2015). Moreover, four SNPs in LD with rs12955474 reached Bonferroni-corrected significance (rs10503028, ß = 0.16, p = 1.0 × 10^−4^; rs34889120, ß = 0.16, p = 1.0 × 10^−4^; rs35781152, ß = 0.16, p = 1.0 × 10^−4^; rs35781152, ß = 0.16, p = 1.2 × 10^−4^).

In gene set enrichment analysis no GO term reached the Bonferroni-corrected level of significance (*p* = 6.8 × 10^−6^) for HT. “Response to interleukin 12 (GO: 0070671)” (*p* = 1.5 × 10^−5^) was the strongest association.

Since there was no significant SNP h^2^for HTA, we did not perform genetic correlation or PRS analyses for HTA. However, since it might benefit future studies interested in HTA, we conducted an explorative GWAS analysis (Supplementary Table S6; Supplementary Figures S8-S11).

#### PRS

PRS were tested for IQ, EA and four neurodevelopmental conditions based on the behavioral correlations between HT and cognitive traits (Figure 2). PRS for ADHD showed an association with HT (Table 1, Supplementary Figure S12), indicating that higher genetic risk for ADHD is associated with higher HT, i.e. worse hearing. In contrast, schizophrenia PRS showed a negative association with HT, suggesting that higher genetic risk for schizophrenia is associated with lower HT, i.e. better hearing (Table 1, Supplementary Figure S13). PRS for EA also reached significance for HT (Table 1, Supplementary Figure S14). The negative association suggests that a genetic liability towards higher EA is associated with better hearing.

**Table 1:**
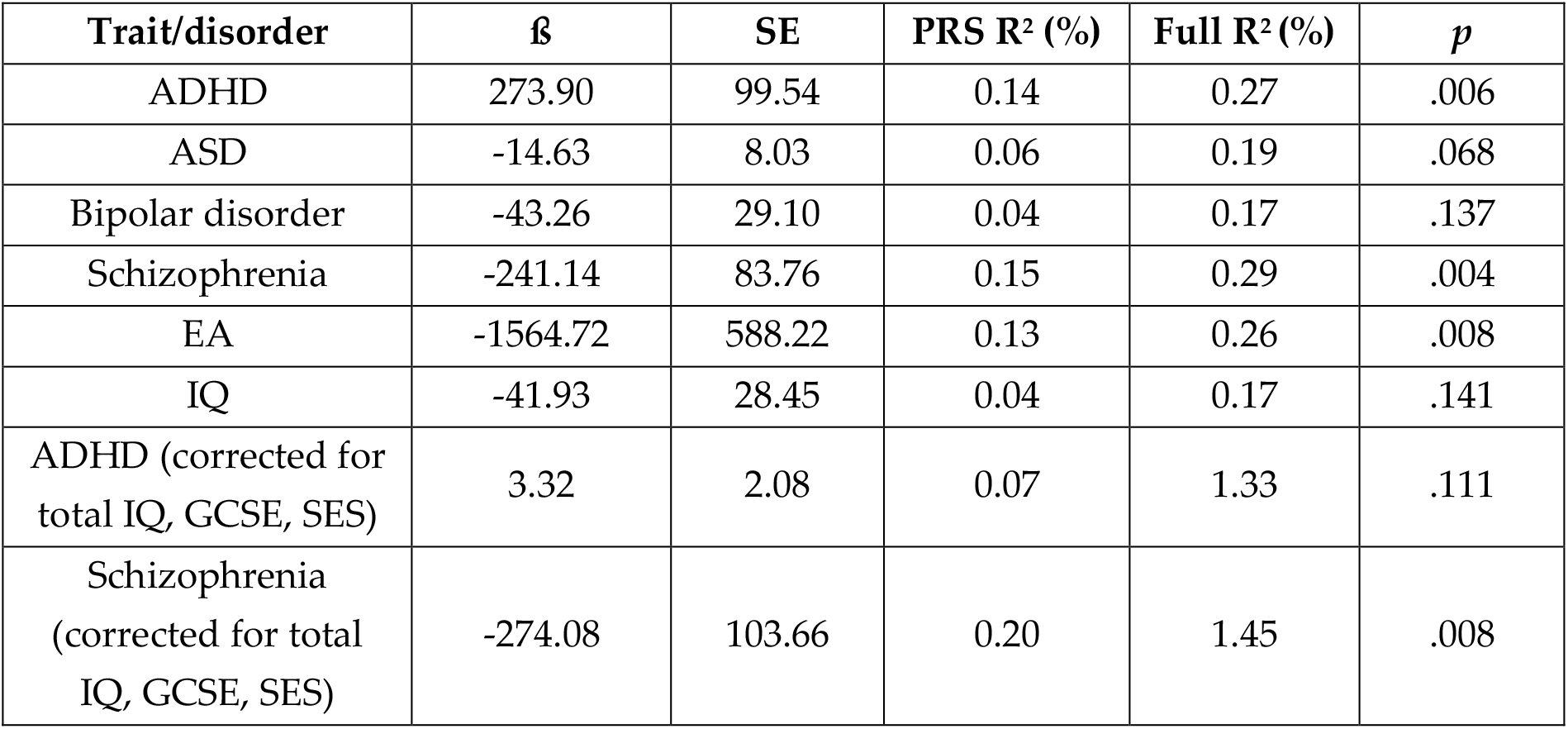
Results of PRS analysis on HT.

We next tested whether the associations between ADHD and schizophrenia PRS with HT were mediated by cognitive skills and SES. We thus reran the PRS analysis on HT with ADHD and schizophrenia as training GWAS and using sex, the first two ancestry-informative principal components, total IQ, GCSE and SES as covariates. The effect of ADHD PRS on HT disappeared after adjusting for total IQ, GCSE and SES (Table 1, Supplementary Figure S15), suggesting that the association is mediated by these variables. In contrast, the effect of schizophrenia PRS on HT remained similar after adjusting for total IQ, GCSE and SES (Table 1, Supplementary Figure S16), suggesting that the association is not mediated by cognitive skills and SES.

## Discussion

We dissect the relationship between hearing measures and cognitive abilities and neurodevelopmental disorders both at the phenotypic and genetic level. We confirm that better hearing is associated with better performance on a range of cognitive tasks (Figure 2). We also report the results of the first GWAS on HT in children. Single marker, gene-based and gene set enrichment analyses did not lead to any statistically significant results. However, PRS for EA and schizophrenia were significantly associated with HT (Table 1), with genetic liability for both traits associated with better hearing.

Phenotypic analysis for hearing measures showed sex-specific developmental effects for both HT and HTA. Our data (*n* = 6,743) show lower HT in boys compared to girls (Figure 1). The sex effect was very small and might only be detectable in reasonably sized samples, explaining why no sex effect was reported in a previous study of *n* = 1869 children and adolescents (Park et al., 2016). In adults, the reverse effect has been reported with better hearing in females compared to males (McFadden, 1993). In our data, HTA revealed an overall left ear advantage with a lower air conduction threshold of 0.28 dB on average (Figure 1), replicating the results from smaller studies in children (Rahko & Karma, 1989; Roche et al., 1978). The effect was driven by females, suggesting that the sex effect on HTA reported in adults (i.e. stronger right ear advantage in males) seems to be established already in children. It is possible that a developmental shift towards the right ear in both sexes is driven by environmental factors, as continuous noise exposure (e.g. industrial noise) has been shown to affect the left ear more than the right (Nageris, Raveh, Zilberberg, & Attias, 2007). A possible limitation of the current study is that stimulus presentation was not randomized, but the right ear was always tested first. Testing the right ear first has been more common in previous studies than vice versa, which could either result in a learning effect (favoring the left ear) or in a fatigue effect (favoring the right ear) (Pirilä, Jounio-Ervasti, & Sorri, 1992). However, in adults, a right ear advantage is more common even in studies in which the order of stimulus presentation has been randomized (Pirilä et al., 1992), so the effect of stimulus presentation should be minimal.

Consistent with previous studies (Moore et al., 2019), we found that in the normal range of variation, HT is negatively associated with several cognitive skills on the behavioral level (Figure 2). Although not significant, we found more negative HTA (i.e. more leftward asymmetry) in neurodevelopmental conditions including ASD compared to controls. Different types of asymmetry such as structural brain asymmetry (Postema et al., 2019), frontal alpha asymmetry (Gabard-Durnam, Tierney, Vogel-Farley, Tager-Flusberg, & Nelson, 2015), language processing (Herringshaw, Ammons, DeRamus, & Kana, 2016) and handedness (Markou, Ahtam, & Papadatou-Pastou, 2017) have been implicated in ASD. Therefore, it might be worth collecting HTA measures more systematically to further investigate the link between asymmetries and disorders.

Because there was no significant SNP h^2^for HTA, downstream genomic analyses were only performed for HT. Although no single marker trait associations reached significance in the GWAS (Supplementary Figure S4), the top marker on chromosome 15 (rs1039444) is located in an intron of *RAB8B*, which encodes for a GTPase that is expressed in inner and outer hair cells and is involved in autosomal recessive deafness (Heidrych et al., 2008). Targeted analysis for markers reported in previous GWAS for HT in adults replicated association with one marker, rs12955474, which is located in an intron of the *CCBE1* gene (Fransen et al., 2015). Other markers in this gene have been associated with depression (Power et al., 2013) and left entorhinal cortex volume (Zhao et al., 2019). The sample size of our study is rather small for general GWAS standard, but of comparable size to GWAS conducted so far for hearing measures in adults. In fact, air conduction thresholds are not routinely collected in large-scale population studies. For example, the UK Biobank includes phenotypic information on hearing ability as self-reported hearing difficulty or use of hearing aids (*n* > 300,000) (Kalra et al., 2019; Wells et al., 2019). Similarly, a GWAS on age-related hearing loss in the Genetic Epidemiology Research on Adult Health and Aging (GERA) cohort (*n* > 50,000) used a case control design, identifying cases based on health records (ICD-9 diagnosis) (Hoffmann et al., 2016). GWAS on quantitative measures of hearing ability were limited to smaller sample sizes below 6,000 subjects for individual samples (Nolan et al., 2013; Vuckovic et al., 2015). Systematic collection of air conduction thresholds in both ears in children would enable larger genetic studies to dissect the links between hearing, cognition and neurodevelopment.

Genetic correlation analysis (Figure 3) revealed associations between HT and listening comprehension, reading ability and GCSE scores. In this analysis, standard errors were quite high, showcasing the need for follow up in larger datasets. We found no genetic correlation between verbal, performance or total IQ and HT. Similarly, PRS derived from large datasets did not identify an association between PRS for IQ (capturing mainly verbal and total IQ, Genç et al. (2020)) and HT, suggesting that behavioral associations are not mediated by shared biological pathways. The analysis of PRS for ADHD and schizophrenia instead support a role of genes implicated in neurodevelopmental disorders contributing to HT. Hearing deficits in ADHD have been reported in terms of speech perception (Fuermaier et al., 2018), but not in air conduction thresholds. In our sample, there was no association between HT and ADHD on the behavioral level, however this analysis was based on a too small sample of children meeting the criteria for ADHD (*n* = 21) to make conclusive results (Supplementary Table S2). Moreover, the effect of ADHD PRS on HT disappeared after adjusting for cognitive skills and SES, suggesting that the effect was mediated by cognitive factors.

The strongest PRS association was observed for schizophrenia PRS and HT, in which higher PRS for schizophrenia were associated with better hearing after adjustment for cognitive skills and SES. Most PRS studies report shared genetic risk for different outcomes, e.g. risk for one disorder tends to increase risk for other disorders (Andlauer et al., 2019) or poor cognitive performance (Gialluisi et al., 2019). Instead, we found that PRS for schizophrenia were associated with better hearing. Similarly, PRS for schizophrenia have recently been associated with better language skills, but not overall school performance (Rajagopal et al., 2020). Therefore, our study expands the range of positive outcomes associated to risk of schizophrenia. The association between better hearing and cognitive skills (including language measures) found on the behavioral level (Figure 2) could be based on shared biological pathways which also increase the risk for schizophrenia. On the phenotypic level, previous research did not detect differences in HT between individuals affected by schizophrenia and controls in small samples (*n* = 87) (Prager & Jeste, 1993). In the future, this association might be worth being investigated more systematically in larger cohorts.

In summary, our results highlight behavioral and genetic overlap between cognitive and sensory domains. We find that PRS for EA and schizophrenia are associated with HT, extending previous research reporting positive outcomes for schizophrenia PRS. Future studies should explore in more detail associations between sensory function and cognitive traits.

## Acknowledgments

We are extremely grateful to all the families who took part in this study, the midwives for their help in recruiting them, and the whole ALSPAC team, which includes interviewers, computer and laboratory technicians, clerical workers, research scientists, volunteers, managers, receptionists and nurses. We are grateful to Veera M. Rajagopal for useful comments to the manuscript.

The UK Medical Research Council and Wellcome (Grant ref: 217065/Z/19/Z) and the University of Bristol provide core support for ALSPAC. GWAS data was generated by Sample Logistics and Genotyping Facilities at Wellcome Sanger Institute and LabCorp (Laboratory Corporation of America) using support from 23andMe. This publication is the work of the authors and SP and JS will serve as guarantors for the analysis of the ALSPAC data presented in this paper. JS is funded by the Deutsche Forschungsgemeinschaft (DFG, German Research Foundation, SCHM 3530/1-1, 418445085). SP is funded by the Royal Society (UF150663). Support to the genetic analysis was provided by the St Andrews Bioinformatics Unit funded by the Wellcome Trust [grant 105621/Z/14/Z].

The authors declare no competing financial interests in relation to the work described.

