## Supplement for "Genome-wide association study and polygenic risk score analysis for hearing measures in children"

**Supplementary methods**

Analysis was conducted in the ALSPAC cohort (Boyd et al., 2013; Fraser et al., 2013).

*Exclusion criteria*

Phenotypic data for both the left and the right ear were available for 7,433 children. Exclusion criteria were sensory impairments ($n$ = 29), bilateral or unilateral hearing impairment (average air conduction threshold between 21 and 40 dBHL inclusive; $n$ = 581), bilateral or unilateral sensorineural hearing loss (average air conduction threshold greater than or equal to 21 dBHL; Type A tympanogram indicating normal middle-ear function; $n$ = 152) or high-frequency hearing loss (air conduction thresholds at 2 kHz 20 dB or less, thresholds at 4 and 8 kHz 25 dBHL or greater; $n$ = 135).

*Cognitive measures*

*Reading ability:* Basic reading was assessed using the basic reading subtest of the Wechsler Objective Reading Dimensions (Rust, Golombok, & Trickey, 1993) at age 7. The task consisted of seven pictures and 48 words for decoding and word reading elements. The task was stopped if the child made six consecutive errors. The final reading score was derived by the number of words the child read correctly and corrected for age in weeks.

*Short term memory:* An adaptation of the Nonword Repetition Test (Gathercole, Willis, Baddeley, & Emslie, 1994) was used to assess short term memory at age 8. This comprised twelve nonsense words, four each of 3, 4 and 5 syllables and conforming to English rules for sound combinations. The child was asked to listen to each word via an audio cassette recorder and then repeat each item. The repetition attempt was scored as correct if there was no phonological deviation from the target form. The number of items correctly repeated was scored for each child and corrected for age in weeks.

*Listening comprehension:* The Wechsler Objective Language Dimensions (WOLD Rust (1996)) was administered at age 8. In the listening comprehension subtest, the child listens to the tester read aloud a paragraph about a picture, which the child is shown. The child then answers questions on what they have heard. The child has to make inferences about what was read to them and answer the questions verbally. The task was discontinued if the child got three consecutive questions incorrect. A sum score was calculated as the sum of the items that the child got correct (ranging from 2-15) and corrected for age in weeks at the time of testing.

*WISC performance IQ:* The WISC-III UK (Wechsler, Golombok, & Rust, 1991) was administered at age 8. The WISC comprises five performance subtests (picture completion, coding, picture arrangement, block design, object assembly). With exception of the coding subtest, short forms of subtests were used. Raw scores were made comparable to full subtest scores by multiplication (i.e. by multiplying by 2 when half the subtest had been administered). Using the WISC manual, age-scaled scores were obtained from the raw scores and a total score was calculated for the WISC performance IQ.

*WISC verbal IQ:* The WISC includes five verbal subtests (information/knowledge, similarities, mental arithmetics, vocabulary, comprehension). Age-scaled scores were obtained as described above and a total score was calculated for the WISC verbal IQ.

*WISC total IQ:* The total WISC IQ score was calculated as the sum of all 10 age-scaled WISC subtests (picture completion, coding, picture arrangement, block design, object assembly, information/knowledge, similarities, mental arithmetics, vocabulary, comprehension).

*Communication skills:* Communication skills were assessed using the children's communication checklist (CCC) (Bishop, 1998) at age 9. The CCC consists of 70 items grouped into 9 subscales (intelligibility and fluency, syntax, appropriate initiation, coherence, stereotyped conversation, use of conversational context, conversational rapport). Sum scores were corrected for age in weeks.

*GCSE scores:* Educational attainment (EA) was measured as capped General Certificate of Secondary Education (GCSE) scores. GCSEs are the main qualification taken at the end of compulsory education in the UK. Capped GCSE scores represent the best eight grades at GCSE. GCSE scores were available for 5,402 children.

For all eight cognitive measures, higher values indicate better performance.

*Subsample assignment*

Group assignment followed the same strategy described previously Scerri et al. (2011). Briefly, from the overall ALSPAC sample ($n$ = 15,443) we excluded individuals with incomplete data on measures used for sample assignment and individuals not reporting white European ethnicity. Next, individuals with a WISC performance IQ below 85 were excluded to remove individuals who may have performed worse on these tests due to restricted cognitive skills. From the remaining sample, individuals were assigned to the groups of reading disability (RD, $n$ = 173), language impairment (LI, $n$ = 184), ADHD ($n$ = 26), comorbid combinations of these disorders (LI + RD, $n$ = 47; LI + ADHD, $n$ = 7, RD + ADHD, $n$ = 5; RD + LI + ADHD, $n$ = 3) or unaffected ($n$ = 3,305) according to the following criteria. Numbers of participants were slightly different from previous publications given updates in the most recent release of the ALSPAC data.

*RD:* Children scoring < -1 SD on tests of age-adjusted single-word reading at 7 and 9 years were assigned to the RD group.

*LI:* An assignment of LI was given if an individual scored positive for at least two of the following four criteria, which target different aspects of language problems:

1. CCC score < -1 SD
2. Nonword Repetition < -1 SD
3. WOLD score < -1 SD
4. positive response on speech/language therapy questionnaire

*ASD:* An assignment to ASD was based on maternal report (*Have you ever been told that your child has autism, Asperger's syndrome or autistic spectrum disorder?*) at the age of 9 years.

*ADHD:* An assignment of ADHD was based on a DAWBA DSM-IV clinical diagnosis.

The sample used in the present study (*n* = 6,743) included 401 affected and 2,745 unaffected individuals. The control group was sex-matched to maintain the same M/F ration of 1.57 observed in the cases.

Individuals assigned to more than one group were included in the "comorbidity" subgroup.

| | ***n*** | ***n* male** | ***n* female** | **ratio male:female** |
|---|---|---|---|---|
| **Unaffected** | 2,071 | 1,264 | 807 | 1.57 |
| **Affected** | 419 | 245 | 156 | 1.57 |
| **LI** | 155 | 82 | 73 | 1.12 |
| **RD** | 141 | 87 | 54 | 1.61 |
| **ASD** | 35 | 19 | 16 | 1.19 |
| **ADHD** | 21 | 20 | < 5 | 20.00 |
| **Comorbidity** | 49 | 37 | 12 | 3.08 |

### Supplementary Figures and Tables

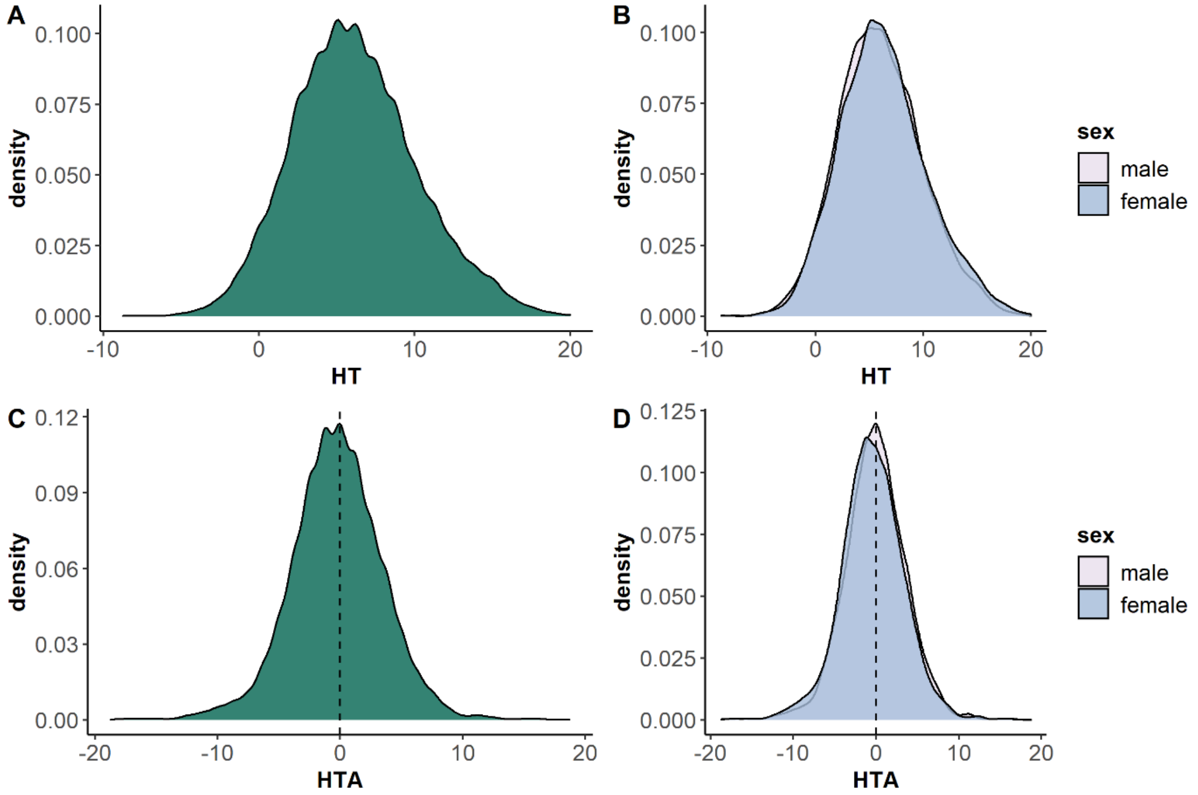

*Figure S1: Distribution of hearing threshold (HT) and hearing threshold asymmetry (HTA). A) Distribution of HT (better ear) in the GWAS subsample (n = 5,344), and B) as a function of sex. C) Distribution of HTA in the GWAS subsample, and D) as a function of sex. The dotted line represents no asymmetry between the left and right ear.*

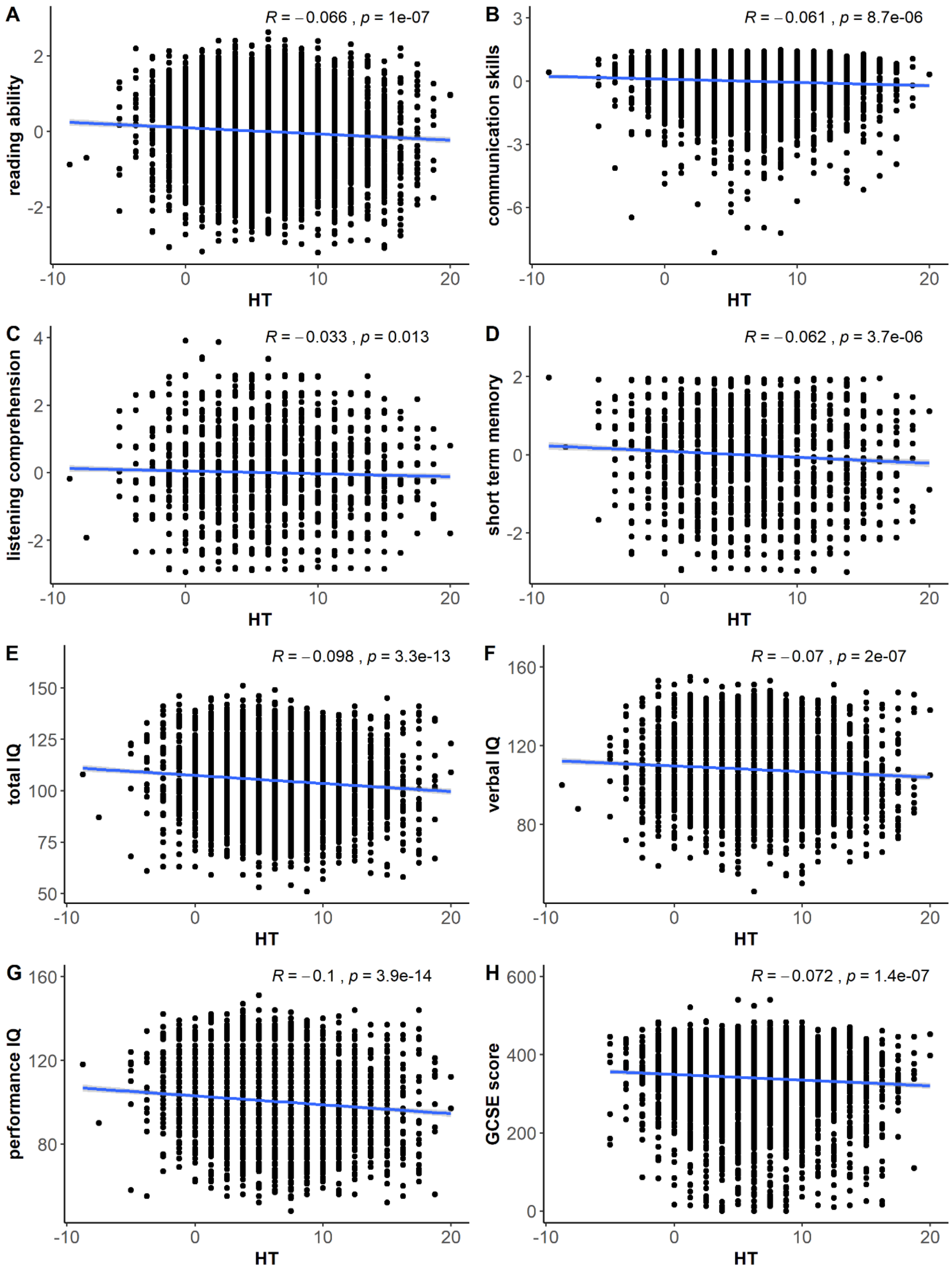

*Figure S2: Correlation plots between cognitive measures and HT. Negative correlation coefficients indicate that lower HT (better hearing) is associated with better cognitive skills.*

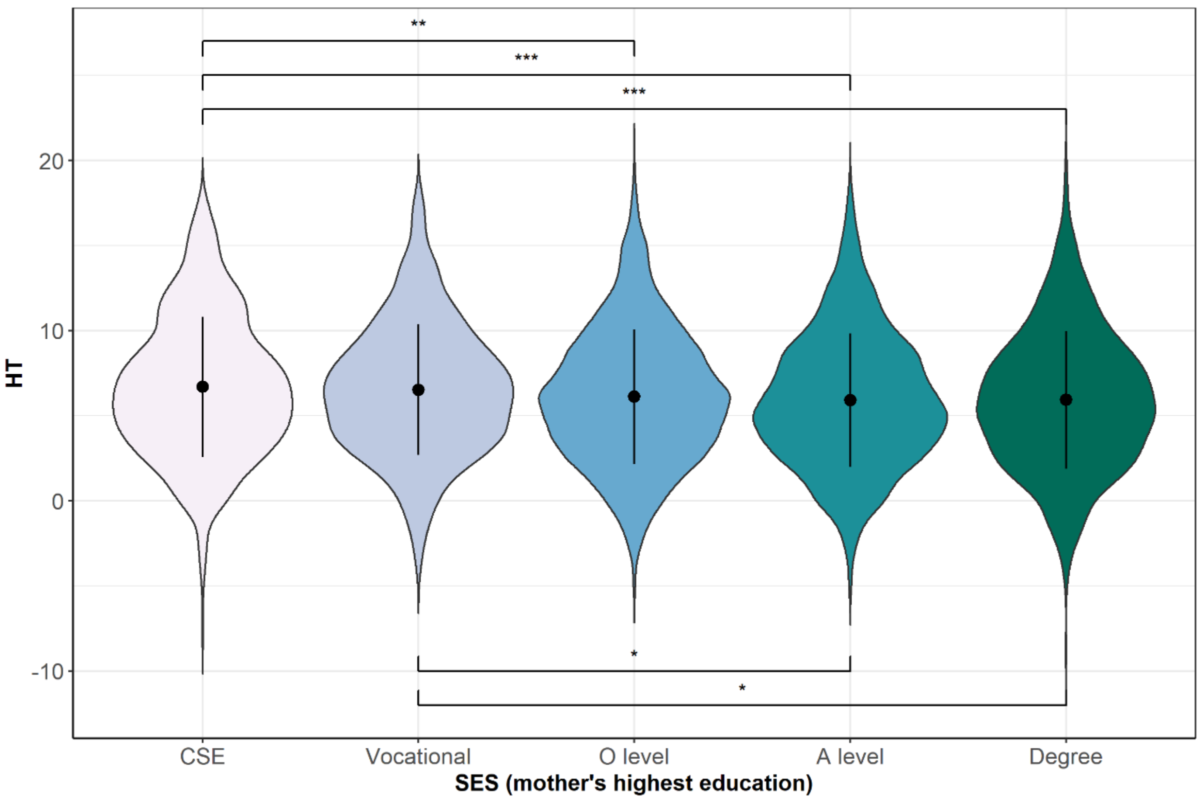

*Figure S3: Effect of SES on HT.* *** $p < .001$, ** $p < .01$, * $p < .05$ *after adjustment for multiple comparisons.*

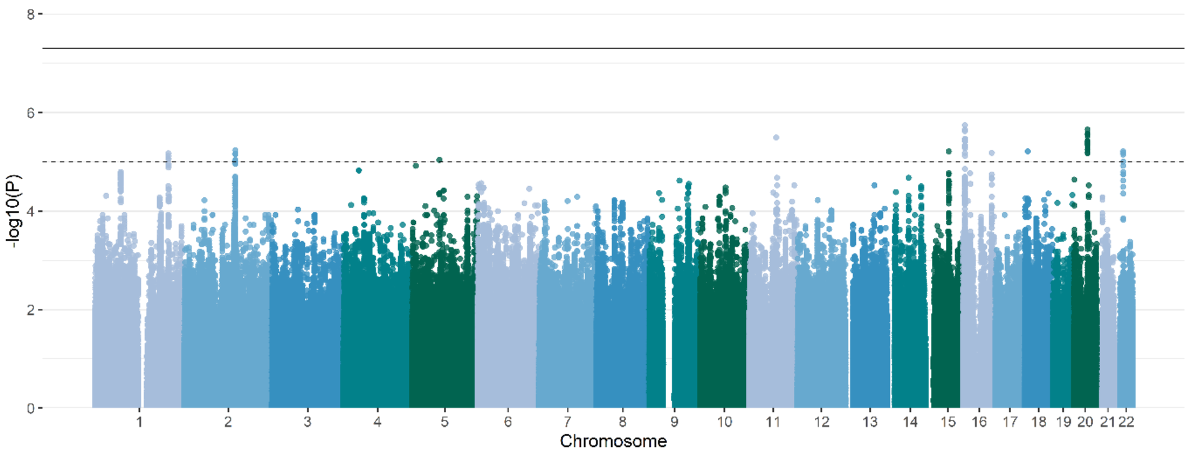

*Figure S4: GWAS Manhattan plot for HT. Association p values are plotted against chromosome and position. The solid line represents the genome-wide significance level ($p = 5 \times 10^{-8}$).*

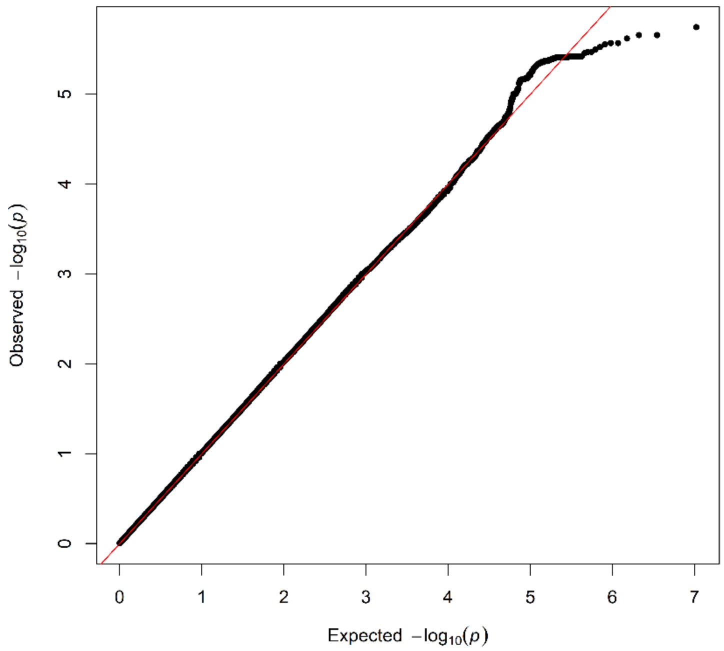

*Figure S5: QQ plot for GWAS on HT.*

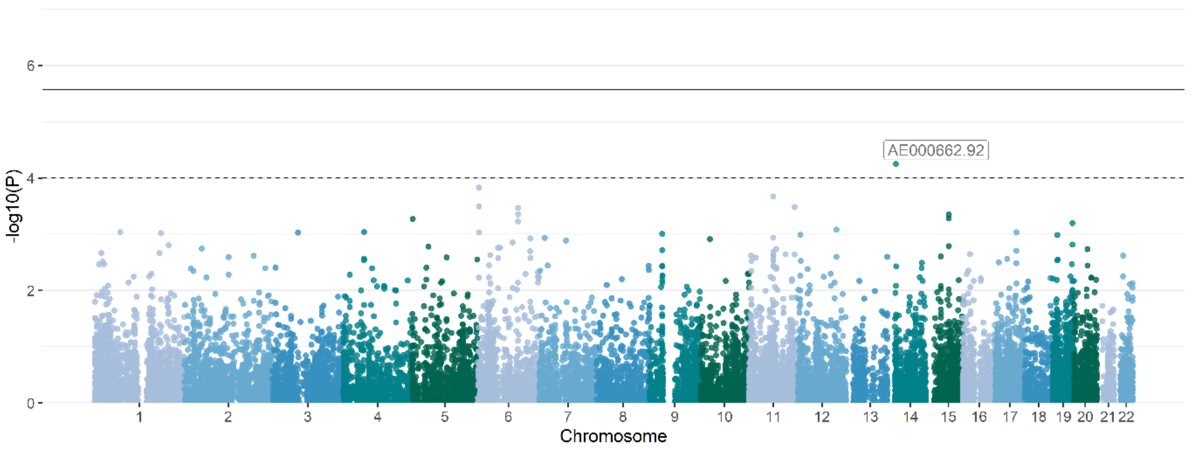

*Figure S6: Gene-based Manhattan plot for HT. Gene-based association p values are plotted against chromosome and position. The solid line represents the genome-wide significance level ($p = 2.7 \times 10^{-6}$).*

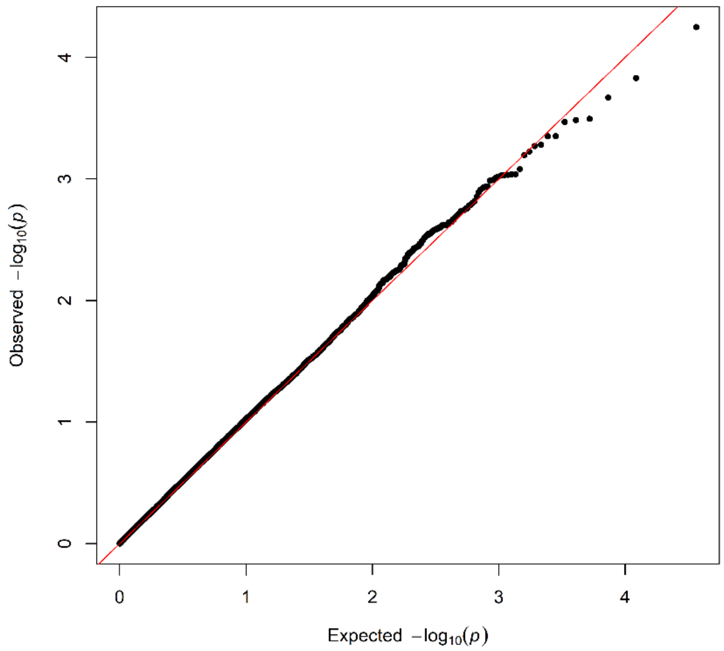

*Figure S7: QQ plot for gene-based GWAS on HT.*

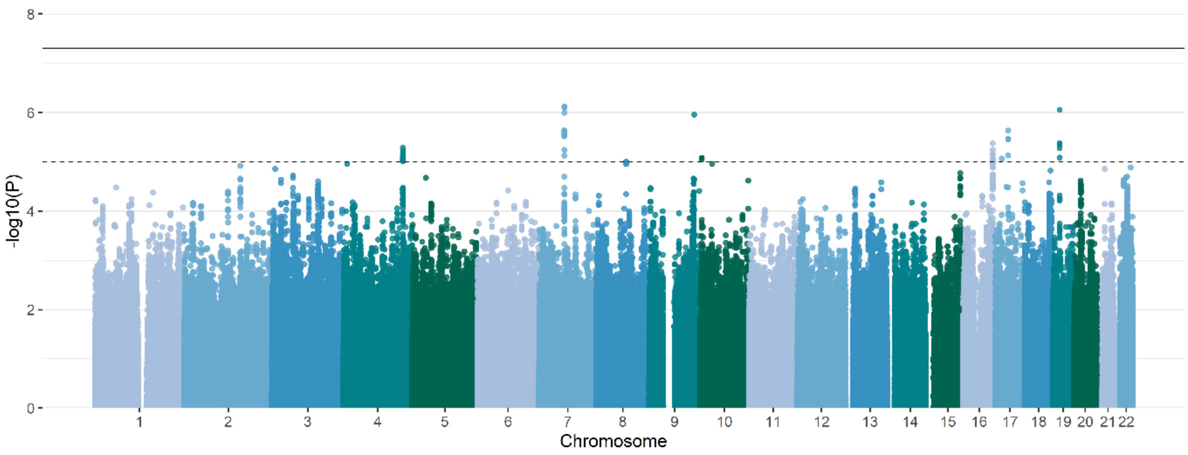

*Figure S8: GWAS Manhattan plot for HTA. Association p values are plotted against chromosome and position. The solid line represents the genome-wide significance level ($p = 5 \times 10^{-8}$).*

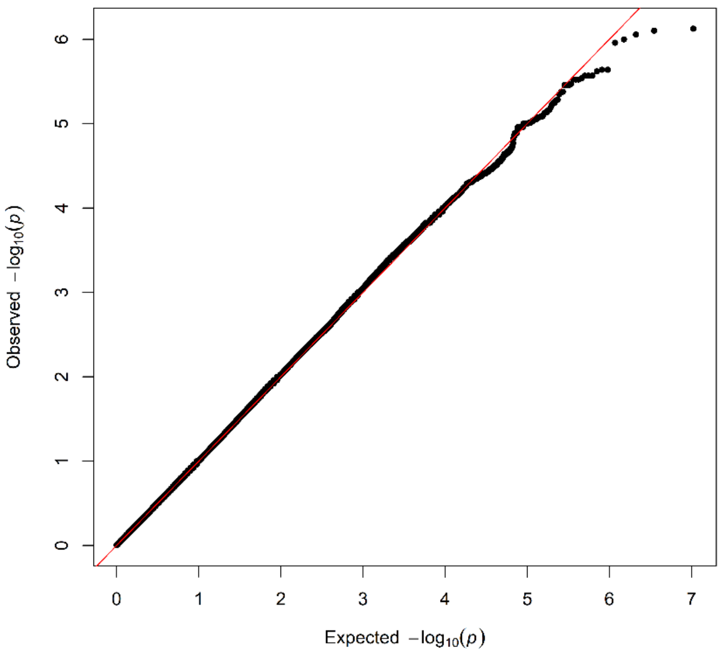

*Figure S9: QQ plot for GWAS on HTA.*

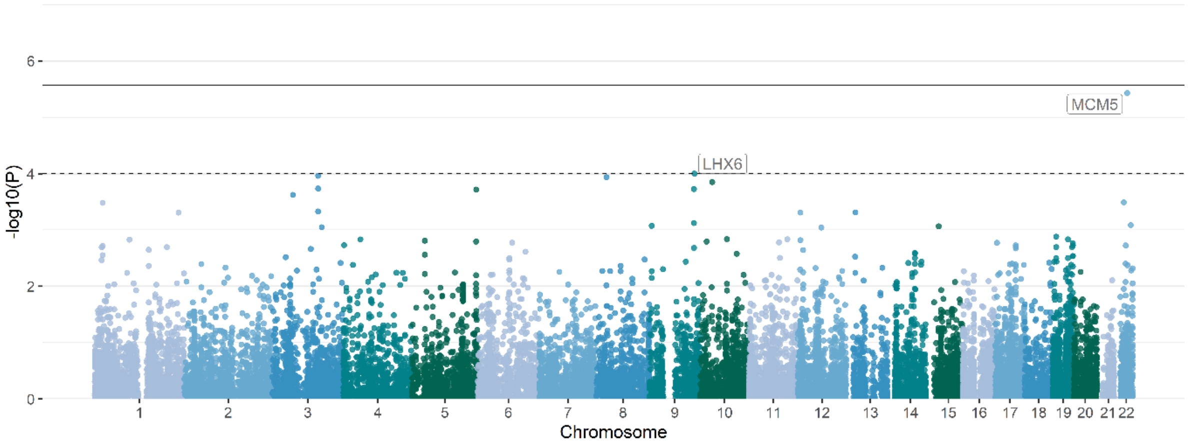

*Figure S10: Gene-based Manhattan plot for HTA. Gene-based association p values are plotted against chromosome and position. The solid line represents the genome-wide significance level ($p = 2.7 \times 10^{-6}$).*

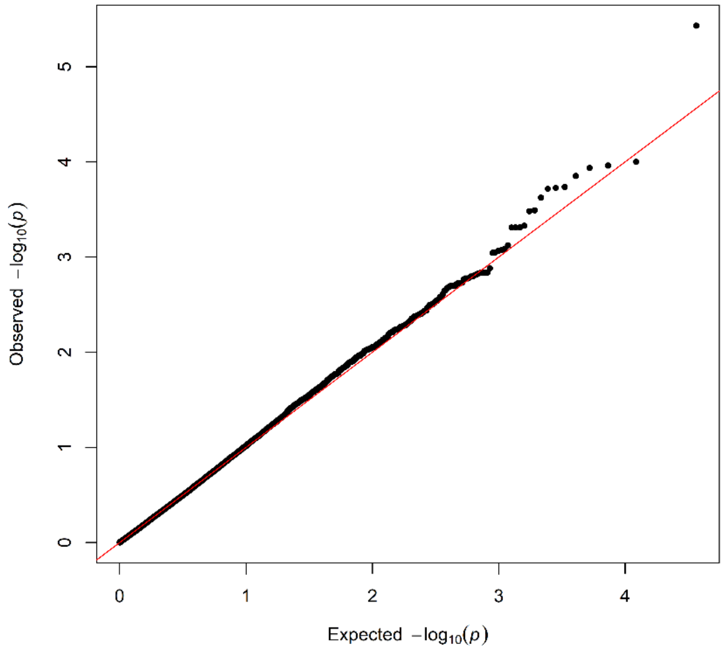

*Figure S11: QQ plot for gene-based GWAS on HTA.*

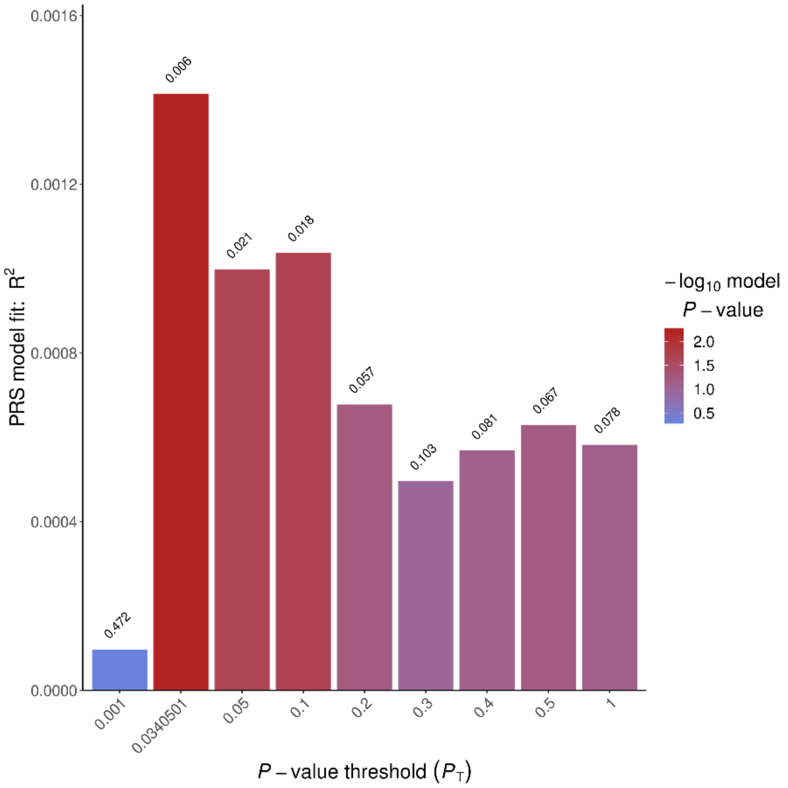

*Figure S12: Results of ADHD PRS analysis. The explained variance in HT by ADHD PRS (PRS $R^2$) is plotted for each training GWAS p value threshold.*

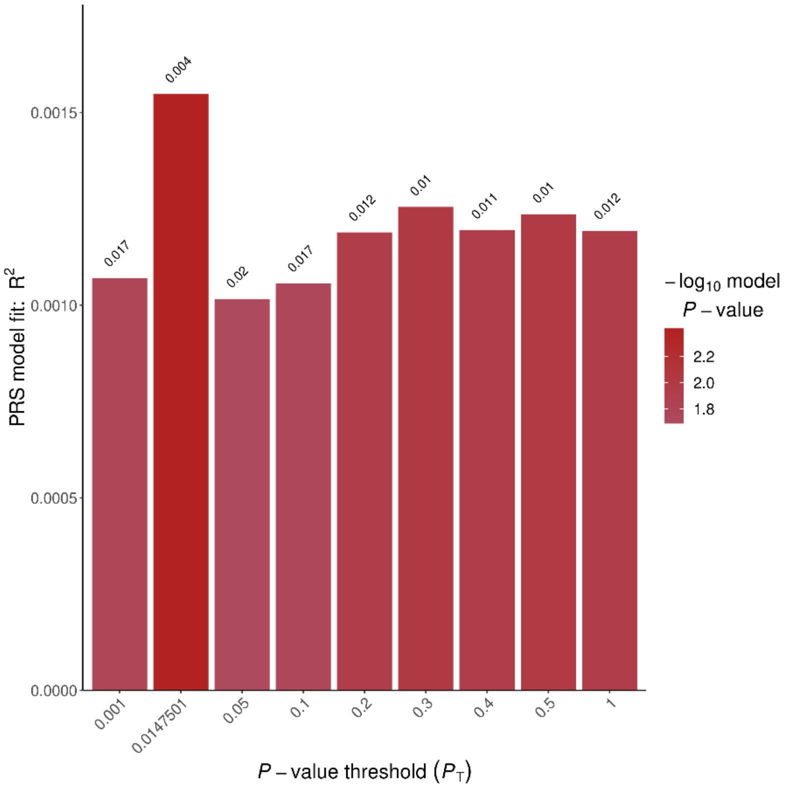

*Figure S13: Results of SCZ PRS analysis. The explained variance in HT by SCZ PRS (PRS $R^2$) is plotted for each training GWAS p value threshold.*

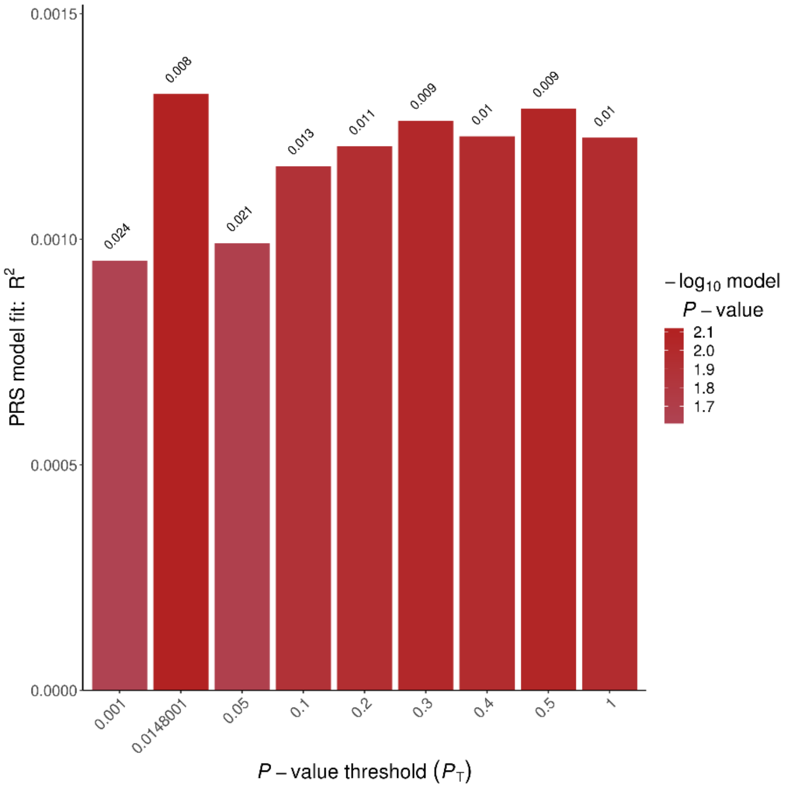

*Figure S14: Results of EA PRS analysis. The explained variance in HT by EA PRS (PRS $R^2$) is plotted for each training GWAS p value threshold.*

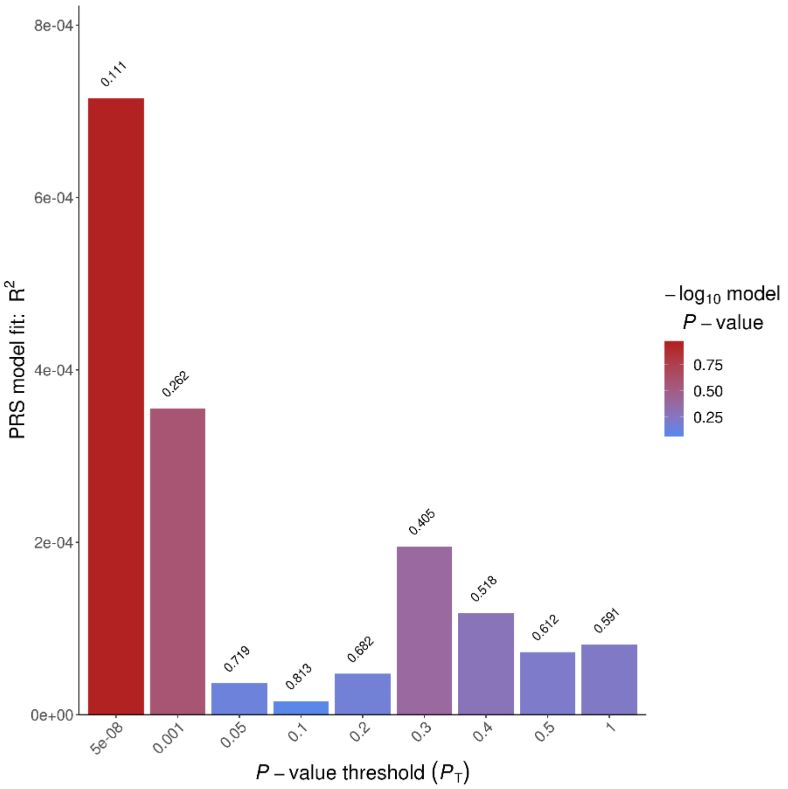

*Figure S15: Results of ADHD PRS analysis including covariates. The explained variance in HT by ADHD PRS (PRS $R^2$) is plotted for each training GWAS p value threshold.*

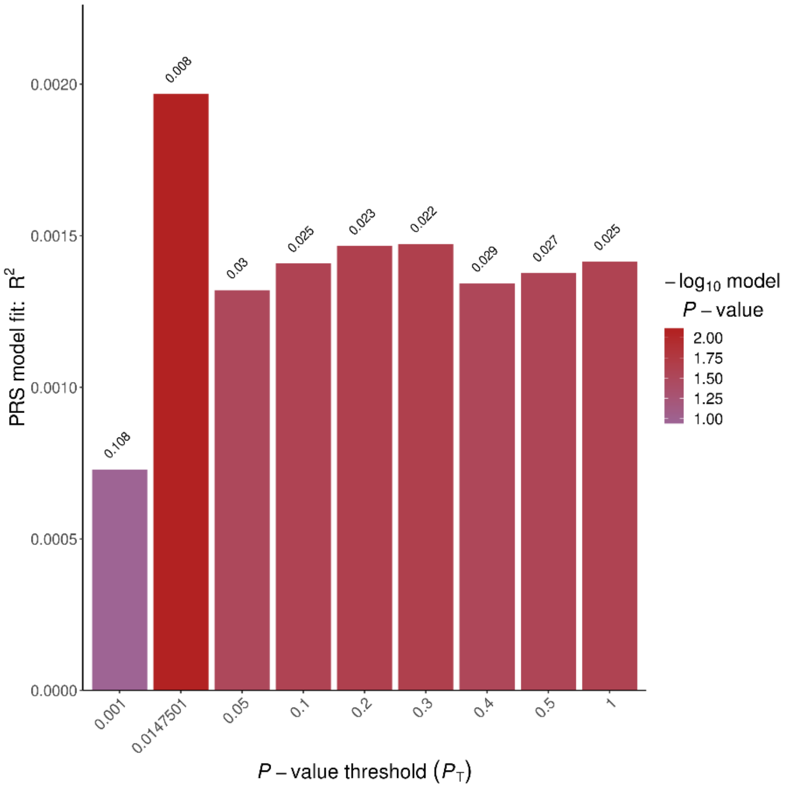

*Figure S16: Results of SCZ PRS analysis including covariates. The explained variance in HT by SCZ PRS (PRS $R^2$) is plotted for each training GWAS p value threshold.*

*Table S1: Optimal p value thresholds of training GWAS (explaining the highest proportion of phenotypic variance in the target sample) and corresponding number of SNPs included in PRS.*

| Training GWAS | *p* value threshold | *n* included SNPs |
|---|---|---|
| ASD | .0001 | 202 |
| ADHD | .0341 | 14,112 |
| BIP | .0013 | 1953 |
| SCZ | .0148 | 12,534 |
| IQ | $5 \times 10^{-8}$ | 261 |
| EA | .0148 | 18,473 |

*Table S2: Results for two-sample t-tests comparing HT between neurodevelopmental conditions and sex-matched controls*

| Condition | *n* | Mean HT | SD HT | *T* | df | *p* |
|---|---|---|---|---|---|---|
| LI | 155 | 6.32 | 3.97 | -1.38 | 177.58 | .170 |
| RD | 141 | 5.83 | 3.81 | 0.10 | 161.21 | .919 |
| ASD | 35 | 6.79 | 4.96 | -1.09 | 34.73 | .283 |
| ADHD | 21 | 5.65 | 4.23 | 0.23 | 20.36 | .821 |
| Comorbidity | 49 | 5.61 | 4.37 | 0.40 | 49.87 | .687 |
| Controls | 2,071 | 5.87 | 3.95 | - | - | - |

*Table S3: Results for two-sample t-tests comparing HTA between neurodevelopmental conditions and sex-matched controls*

| Condition | *n* | Mean HTA | SD HTA | *T* | df | *p* |
|---|---|---|---|---|---|---|
| LI | 155 | -0.47 | 3.80 | 0.71 | 175.91 | .481 |
| RD | 141 | -0.52 | 3.89 | 0.83 | 157.23 | .410 |
| ASD | 35 | -0.86 | 3.80 | 0.95 | 35.07 | .350 |
| ADHD | 21 | -0.89 | 3.07 | 0.96 | 20.58 | .347 |
| Comorbidity | 49 | -0.43 | 3.01 | 0.43 | 51.39 | .668 |
| Controls | 2,071 | -0.24 | 3.65 | - | - | - |

*Table S4: Top SNPs (association p < .00001) for HT GWAS.*

| rsID | Chr:pos | Non-effect allele | Effect allele | Frequency of effect allele | $\chi^2$ | ß | SE ß | *p* | Nearest gene upstream | Distance to nearest gene upstream (bp) | Nearest gene downstream | Distance to nearest gene downstream (bp) | Function |
|---|---|---|---|---|---|---|---|---|---|---|---|---|---|
| rs11644235 | 16:5601755 | G | T | 0.58 | 22.77 | 0.09 | 0.02 | 1.80E-06 | RP11-420N3.2 | 0 | RP11-420N3.2 | 0 | ncRNA_intronic |
| rs11643126 | 16:5601731 | G | A | 0.58 | 22.40 | 0.09 | 0.02 | 2.20E-06 | RP11-420N3.2 | 0 | RP11-420N3.2 | 0 | ncRNA_intronic |
| rs2757517 | 20:37375385 | T | C | 0.64 | 22.45 | 0.09 | 0.02 | 2.20E-06 | SLC32A1 | 17370 | ACTR5 | 1700 | intergenic |
| rs34556976 | 16:5601181 | C | T | 0.58 | 22.23 | 0.09 | 0.02 | 2.40E-06 | RP11-420N3.2 | 0 | RP11-420N3.2 | 0 | ncRNA_intronic |
| rs2748657 | 20:37374016 | A | G | 0.64 | 22.05 | 0.09 | 0.02 | 2.70E-06 | SLC32A1 | 16001 | ACTR5 | 3069 | intergenic |
| rs6027727 | 20:37375469 | T | G | 0.64 | 21.99 | 0.09 | 0.02 | 2.70E-06 | SLC32A1 | 17454 | ACTR5 | 1616 | intergenic |
| rs2748659 | 20:37375155 | G | A | 0.64 | 21.96 | 0.09 | 0.02 | 2.80E-06 | SLC32A1 | 17140 | ACTR5 | 1930 | intergenic |
| rs2748658 | 20:37375140 | C | T | 0.64 | 21.81 | 0.09 | 0.02 | 3.00E-06 | SLC32A1 | 17125 | ACTR5 | 1945 | intergenic |
| rs60995418 | 11:75882890 | A | G | 0.58 | 21.68 | -0.09 | 0.02 | 3.20E-06 | UVRAG | 28651 | WNT11 | 14479 | intergenic |
| rs7200483 | 16:5602094 | G | C | 0.58 | 21.55 | 0.09 | 0.02 | 3.40E-06 | RP11-420N3.2 | 0 | RP11-420N3.2 | 0 | ncRNA_intronic |
| rs6500696 | 16:5602316 | A | G | 0.58 | 21.55 | 0.09 | 0.02 | 3.40E-06 | RP11-420N3.2 | 0 | RP11-420N3.2 | 0 | ncRNA_intronic |
| rs6500697 | 16:5602448 | C | T | 0.58 | 21.54 | 0.09 | 0.02 | 3.50E-06 | RP11-420N3.2 | 0 | RP11-420N3.2 | 0 | ncRNA_intronic |
| rs34926967 | 16:5601948 | A | G | 0.58 | 21.38 | 0.09 | 0.02 | 3.80E-06 | RP11-420N3.2 | 0 | RP11-420N3.2 | 0 | ncRNA_intronic |
| rs60695748 | 16:5602835 | C | T | 0.58 | 21.35 | 0.09 | 0.02 | 3.80E-06 | RP11-420N3.2 | 0 | RP11-420N3.2 | 0 | ncRNA_intronic |
| rs34788411 | 16:5602954 | G | A | 0.58 | 21.35 | 0.09 | 0.02 | 3.80E-06 | RP11-420N3.2 | 0 | RP11-420N3.2 | 0 | ncRNA_intronic |
| rs57902618 | 16:5603024 | A | G | 0.58 | 21.37 | 0.09 | 0.02 | 3.80E-06 | RP11-420N3.2 | 0 | RP11-420N3.2 | 0 | ncRNA_intronic |
| rs2250596 | 20:37371485 | C | T | 0.64 | 21.35 | 0.09 | 0.02 | 3.80E-06 | SLC32A1 | 13470 | ACTR5 | 5600 | intergenic |
| rs2748654 | 20:37371770 | A | G | 0.64 | 21.34 | 0.09 | 0.02 | 3.80E-06 | SLC32A1 | 13755 | ACTR5 | 5315 | intergenic |
| rs2250447 | 20:37370144 | G | C | 0.64 | 21.31 | 0.09 | 0.02 | 3.90E-06 | SLC32A1 | 12129 | ACTR5 | 6941 | intergenic |
| rs2250455 | 20:37370218 | C | T | 0.64 | 21.30 | 0.09 | 0.02 | 3.90E-06 | SLC32A1 | 12203 | ACTR5 | 6867 | intergenic |
| rs2250474 | 20:37370933 | G | A | 0.64 | 21.31 | 0.09 | 0.02 | 3.90E-06 | SLC32A1 | 12918 | ACTR5 | 6152 | intergenic |
| rs2425365 | 20:37372068 | C | T | 0.64 | 21.31 | 0.09 | 0.02 | 3.90E-06 | SLC32A1 | 14053 | ACTR5 | 5017 | intergenic |
| rs2748655 | 20:37372364 | C | T | 0.64 | 21.32 | 0.09 | 0.02 | 3.90E-06 | SLC32A1 | 14349 | ACTR5 | 4721 | intergenic |
| rs6027723 | 20:37372547 | G | T | 0.64 | 21.32 | 0.09 | 0.02 | 3.90E-06 | SLC32A1 | 14532 | ACTR5 | 4538 | intergenic |
| rs2748656 | 20:37372674 | G | A | 0.64 | 21.33 | 0.09 | 0.02 | 3.90E-06 | SLC32A1 | 14659 | ACTR5 | 4411 | intergenic |
| rs2250741 | 20:37372810 | A | G | 0.64 | 21.33 | 0.09 | 0.02 | 3.90E-06 | SLC32A1 | 14795 | ACTR5 | 4275 | intergenic |
| rs2144537 | 20:37369150 | T | C | 0.64 | 21.24 | 0.09 | 0.02 | 4.00E-06 | SLC32A1 | 11135 | ACTR5 | 7935 | intergenic |
| rs2025155 | 20:37371684 | G | A | 0.36 | 21.26 | -0.09 | 0.02 | 4.00E-06 | SLC32A1 | 13669 | ACTR5 | 5401 | intergenic |
| rs2263749 | 20:37368708 | A | C | 0.64 | 21.20 | 0.09 | 0.02 | 4.10E-06 | SLC32A1 | 10693 | ACTR5 | 8377 | intergenic |

| | | | | | | | | | | | | | |
|---|---|---|---|---|---|---|---|---|---|---|---|---|---|
| rs2250240 | 20:37368743 | A | G | 0.64 | 21.21 | 0.09 | 0.02 | 4.10E-06 | SLC32A1 | 10728 | ACTR5 | 8342 | intergenic |
| rs2144539 | 20:37368249 | A | G | 0.64 | 21.18 | 0.09 | 0.02 | 4.20E-06 | SLC32A1 | 10234 | ACTR5 | 8836 | intergenic |
| rs2144538 | 20:37368269 | T | C | 0.64 | 21.18 | 0.09 | 0.02 | 4.20E-06 | SLC32A1 | 10254 | ACTR5 | 8816 | intergenic |
| rs2748649 | 20:37366931 | C | T | 0.64 | 21.11 | 0.09 | 0.02 | 4.30E-06 | SLC32A1 | 8916 | ACTR5 | 10154 | intergenic |
| rs2748650 | 20:37367001 | A | G | 0.64 | 21.12 | 0.09 | 0.02 | 4.30E-06 | SLC32A1 | 8986 | ACTR5 | 10084 | intergenic |
| rs2748651 | 20:37367456 | G | A | 0.64 | 21.11 | 0.09 | 0.02 | 4.30E-06 | SLC32A1 | 9441 | ACTR5 | 9629 | intergenic |
| rs2748652 | 20:37367503 | C | T | 0.64 | 21.11 | 0.09 | 0.02 | 4.30E-06 | SLC32A1 | 9488 | ACTR5 | 9582 | intergenic |
| rs6100973 | 20:37366570 | T | G | 0.64 | 21.07 | 0.09 | 0.02 | 4.40E-06 | SLC32A1 | 8555 | ACTR5 | 10515 | intergenic |
| rs6100974 | 20:37366591 | A | G | 0.64 | 21.08 | 0.09 | 0.02 | 4.40E-06 | SLC32A1 | 8576 | ACTR5 | 10494 | intergenic |
| rs6128859 | 20:37366807 | T | G | 0.64 | 21.09 | 0.09 | 0.02 | 4.40E-06 | SLC32A1 | 8792 | ACTR5 | 10278 | intergenic |
| rs2475387 | 20:37365330 | A | G | 0.64 | 21.05 | 0.09 | 0.02 | 4.50E-06 | SLC32A1 | 7315 | ACTR5 | 11755 | intergenic |
| rs2263748 | 20:37366096 | C | A | 0.64 | 21.05 | 0.09 | 0.02 | 4.50E-06 | SLC32A1 | 8081 | ACTR5 | 10989 | intergenic |
| rs729172 | 16:5606197 | T | G | 0.58 | 20.98 | 0.09 | 0.02 | 4.60E-06 | RP11-420N3.2 | 0 | | | ncRNA_intronic |
| rs6064943 | 20:37365424 | T | A | 0.64 | 21.00 | 0.09 | 0.02 | 4.60E-06 | SLC32A1 | 7409 | ACTR5 | 11661 | intergenic |
| rs11640110 | 16:5606416 | C | T | 0.58 | 20.97 | 0.09 | 0.02 | 4.70E-06 | RP11-420N3.2 | 0 | RP11-420N3.2 | 0 | ncRNA_intronic |
| rs11645142 | 16:5606879 | T | C | 0.58 | 20.90 | 0.09 | 0.02 | 4.80E-06 | RP11-420N3.2 | 0 | RP11-420N3.2 | 0 | ncRNA_intronic |
| rs11644922 | 16:5607099 | A | G | 0.58 | 20.86 | 0.09 | 0.02 | 4.90E-06 | RP11-420N3.2 | 0 | RP11-420N3.2 | 0 | ncRNA_intronic |
| rs11640221 | 16:5606843 | G | A | 0.58 | 20.83 | 0.09 | 0.02 | 5.00E-06 | RP11-420N3.2 | 0 | RP11-420N3.2 | 0 | ncRNA_intronic |
| rs2475384 | 20:37365129 | G | A | 0.64 | 20.80 | 0.09 | 0.02 | 5.10E-06 | SLC32A1 | 7114 | ACTR5 | 11956 | intergenic |
| rs11644816 | 16:5606854 | C | G | 0.58 | 20.72 | 0.09 | 0.02 | 5.30E-06 | RP11-420N3.2 | 0 | | | ncRNA_intronic |
| rs2475385 | 20:37365189 | T | C | 0.64 | 20.63 | 0.09 | 0.02 | 5.60E-06 | SLC32A1 | 7174 | ACTR5 | 11896 | intergenic |
| rs2475386 | 20:37365268 | T | C | 0.64 | 20.63 | 0.09 | 0.02 | 5.60E-06 | SLC32A1 | 7253 | ACTR5 | 11817 | intergenic |
| rs1486964 | 2:140973960 | A | T | 0.68 | 20.55 | -0.09 | 0.02 | 5.80E-06 | AC092156.3 | 0 | AC092156.3 | 0 | downstream |
| rs1039444 | 15:63483202 | C | T | 0.88 | 20.44 | 0.13 | 0.03 | 6.10E-06 | RAB8B | 0 | RAB8B | 0 | intronic |
| rs62086586 | 18:8786700 | A | G | 0.86 | 20.44 | -0.13 | 0.03 | 6.10E-06 | SOGA2 | 0 | SOGA2 | 0 | intronic |
| rs62231150 | 22:25248274 | A | G | 0.93 | 20.45 | 0.17 | 0.04 | 6.10E-06 | SGSM1 | 0 | SGSM1 | 0 | intronic |
| rs2180224 | 20:37361043 | T | C | 0.63 | 20.35 | 0.09 | 0.02 | 6.40E-06 | SLC32A1 | 3028 | ACTR5 | 16042 | intergenic |
| rs11641512 | 16:5607313 | C | T | 0.58 | 20.32 | 0.09 | 0.02 | 6.60E-06 | RP11-420N3.2 | 0 | RP11-420N3.2 | 0 | ncRNA_intronic |
| rs4307960 | 16:79879453 | G | A | 0.90 | 20.31 | 0.15 | 0.03 | 6.60E-06 | RP11-345M22.3 | 18406 | RP11-148M9.1 | 195106 | intergenic |
| rs111366960 | 22:25249786 | A | G | 0.93 | 20.30 | 0.17 | 0.04 | 6.60E-06 | SGSM1 | 0 | SGSM1 | 0 | intronic |
| rs6593890 | 1:204647493 | A | G | 0.79 | 20.26 | 0.11 | 0.02 | 6.70E-06 | LRRN2 | 0 | LRRN2 | 0 | intronic |
| rs1010321 | 20:37360944 | C | A | 0.63 | 20.29 | 0.09 | 0.02 | 6.70E-06 | SLC32A1 | 2929 | ACTR5 | 16141 | intergenic |
| rs1047874 | 2:140989843 | G | A | 0.34 | 20.25 | 0.09 | 0.02 | 6.80E-06 | LRP1B | 0 | LRP1B | 0 | UTR3 |
| rs1486958 | 2:140993750 | T | G | 0.34 | 20.25 | 0.09 | 0.02 | 6.80E-06 | LRP1B | 0 | LRP1B | 0 | intronic |
| rs13416149 | 2:140993930 | C | G | 0.34 | 20.24 | 0.09 | 0.02 | 6.80E-06 | LRP1B | 0 | LRP1B | 0 | intronic |

| | | | | | | | | | | | | | |
|---|---|---|---|---|---|---|---|---|---|---|---|---|---|
| rs1486966 | 2:140979489 | A | T | 0.68 | 20.21 | -0.09 | 0.02 | 6.90E-06 | MTND1P27 | 0 | MTND1P27 | 0 | upstream |
| rs1486968 | 2:140979600 | A | G | 0.68 | 20.21 | -0.09 | 0.02 | 6.90E-06 | MTND1P27 | 0 | MTND1P27 | 0 | upstream |
| rs13007053 | 2:140986237 | C | T | 0.34 | 20.21 | 0.09 | 0.02 | 6.90E-06 | MTND1P27 | 7183 | LRP1B | 2755 | intergenic |
| rs1492388 | 2:140995095 | G | T | 0.34 | 20.23 | 0.09 | 0.02 | 6.90E-06 | LRP1B | 0 | LRP1B | 0 | intronic |
| rs2046562 | 2:140984124 | A | G | 0.34 | 20.18 | 0.09 | 0.02 | 7.00E-06 | MTND1P27 | 5070 | LRP1B | 4868 | intergenic |
| rs13020859 | 2:140988151 | C | T | 0.34 | 20.18 | 0.09 | 0.02 | 7.00E-06 | LRP1B | 0 | LRP1B | 0 | downstream |
| rs62231152 | 22:25259433 | A | G | 0.93 | 20.17 | 0.17 | 0.04 | 7.10E-06 | SGSM1 | 0 | SGSM1 | 0 | intronic |
| rs55636403 | 1:204648046 | C | A | 0.79 | 20.03 | 0.11 | 0.02 | 7.60E-06 | LRRN2 | 0 | LRRN2 | 0 | intronic |
| rs34707192 | 16:5596517 | C | A | 0.55 | 20.02 | 0.09 | 0.02 | 7.60E-06 | RP11-420N3.2 | 0 | RP11-420N3.2 | 0 | ncRNA_intronic |
| rs10900420 | 1:204646261 | A | G | 0.79 | 19.77 | 0.11 | 0.02 | 8.70E-06 | LRRN2 | 0 | LRRN2 | 0 | intronic |
| rs10900421 | 1:204646295 | A | G | 0.79 | 19.77 | 0.11 | 0.02 | 8.70E-06 | LRRN2 | 0 | LRRN2 | 0 | intronic |
| rs11686351 | 2:140984422 | C | T | 0.32 | 19.68 | 0.09 | 0.02 | 9.10E-06 | MTND1P27 | 5368 | LRP1B | 4570 | intergenic |
| rs7448529 | 5:75494996 | C | A | 0.45 | 19.70 | -0.09 | 0.02 | 9.10E-06 | SV2C | 0 | SV2C | 0 | intronic |
| rs2046566 | 2:140998776 | T | G | 0.32 | 19.62 | 0.09 | 0.02 | 9.50E-06 | LRP1B | 0 | LRP1B | 0 | intronic |
| rs2046981 | 2:140984797 | A | T | 0.32 | 19.60 | 0.09 | 0.02 | 9.60E-06 | MTND1P27 | 5743 | LRP1B | 4195 | intergenic |
| rs12485138 | 22:25238806 | A | G | 0.93 | 19.53 | 0.17 | 0.04 | 9.90E-06 | SGSM1 | 0 | SGSM1 | 0 | intronic |

*Table S5: Results of replication analysis for HT.*

| | | Current study | | | | | | | | | Previous studies | | | |
|---|---|---|---|---|---|---|---|---|---|---|---|---|---|---|
| **rsID** | **Chr:pos** | **Non-effect allele** | **Effect allele** | **Frequency of effect allele** | **ß** | **SE ß** | ***p*** | **Nearest gene** | **Distance to nearest gene (bp)** | **Function** | **ß** | ***p*** | **Phenotype** | **Study** |
| rs12955474 | 18:57152038 | C | T | 0.06 | 0.17 | 0.04 | 8.10E-05 | CCBE1 | 0 | intronic | -0.10 | 3.57E-07 | PC3 | (Fransen et al., 2015) |
| rs10503028 | 18:57154357 | A | C | 0.06 | 0.16 | 0.04 | 1.00E-04 | CCBE1 | 0 | intronic | | | | |
| rs34889120 | 18:57157675 | T | G | 0.08 | 0.16 | 0.04 | 1.00E-04 | CCBE1 | 0 | intronic | | | | |
| rs35781152 | 18:57159095 | G | C | 0.06 | 0.16 | 0.04 | 1.00E-04 | CCBE1 | 0 | intronic | | | | |
| rs1557402 | 18:57159558 | C | T | 0.07 | 0.16 | 0.04 | 0.00012 | CCBE1 | 0 | intronic | | | | |

*Table S6: Top SNPs (association p < .00001) for HTA GWAS.*

| rsID | Chr:pos | Non-effect allele | Effect allele | Frequency of effect allele | $\chi^2$ | ß | SE ß | *p* | Nearest gene upstream | Distance to nearest gene upstream (bp) | Nearest gene downstream | Distance to nearest gene downstream (bp) | Function |
|---|---|---|---|---|---|---|---|---|---|---|---|---|---|
| rs10434985 | 7:70384937 | C | T | 0.79 | 24.49 | 0.12 | 0.02 | 7.50E-07 | RP11-575M4.1 | 81938 | WBSCR17 | 212218 | intergenic |
| rs10434958 | 7:70385174 | C | G | 0.79 | 24.39 | 0.12 | 0.02 | 7.90E-07 | RP11-575M4.1 | 82175 | WBSCR17 | 211981 | intergenic |
| rs4808891 | 19:19084684 | T | C | 0.62 | 24.17 | 0.10 | 0.02 | 8.80E-07 | RN7SL70P | 15617 | SUGP2 | 17013 | intergenic |
| rs11981945 | 7:70385106 | T | C | 0.21 | 23.87 | -0.11 | 0.02 | 1.00E-06 | RP11-575M4.1 | 82107 | WBSCR17 | 212049 | intergenic |
| rs72767793 | 9:124981562 | G | A | 0.72 | 23.77 | -0.10 | 0.02 | 1.10E-06 | LHX6 | 0 | LHX6 | 0 | intronic |
| rs73351943 | 7:70386585 | T | G | 0.79 | 22.32 | 0.11 | 0.02 | 2.30E-06 | RP11-575M4.1 | 83586 | WBSCR17 | 210570 | intergenic |
| rs13380817 | 17:35243712 | A | G | 0.47 | 22.33 | -0.09 | 0.02 | 2.30E-06 | RP11-445F12.1 | 0 | RP11-445F12.1 | 0 | ncRNA_intronic |
| rs1880579 | 7:70390228 | T | C | 0.79 | 22.27 | 0.11 | 0.02 | 2.40E-06 | RP11-575M4.1 | 87229 | WBSCR17 | 206927 | intergenic |
| rs56769182 | 7:70386891 | G | A | 0.79 | 22.02 | 0.11 | 0.02 | 2.70E-06 | RP11-575M4.1 | 83892 | WBSCR17 | 210264 | intergenic |
| rs59926049 | 7:70387084 | T | C | 0.79 | 21.99 | 0.11 | 0.02 | 2.70E-06 | RP11-575M4.1 | 84085 | WBSCR17 | 210071 | intergenic |
| rs61502348 | 7:70387158 | G | A | 0.79 | 22.02 | 0.11 | 0.02 | 2.70E-06 | RP11-575M4.1 | 84159 | WBSCR17 | 209997 | intergenic |
| rs57215230 | 7:70392095 | T | C | 0.79 | 21.91 | 0.11 | 0.02 | 2.90E-06 | RP11-575M4.1 | 89096 | WBSCR17 | 205060 | intergenic |
| rs940728 | 7:70387953 | G | A | 0.79 | 21.82 | 0.11 | 0.02 | 3.00E-06 | RP11-575M4.1 | 84954 | WBSCR17 | 209202 | intergenic |
| rs2103170 | 7:70388178 | G | C | 0.79 | 21.83 | 0.11 | 0.02 | 3.00E-06 | RP11-575M4.1 | 85179 | WBSCR17 | 208977 | intergenic |
| rs58492152 | 7:70393496 | A | G | 0.79 | 21.83 | 0.11 | 0.02 | 3.00E-06 | RP11-575M4.1 | 90497 | WBSCR17 | 203659 | intergenic |
| rs11652230 | 17:35243500 | A | G | 0.47 | 21.56 | -0.09 | 0.02 | 3.40E-06 | RP11-445F12.1 | 0 | RP11-445F12.1 | 0 | ncRNA_intronic |
| rs58249358 | 17:35243896 | C | T | 0.47 | 21.52 | -0.09 | 0.02 | 3.50E-06 | RP11-445F12.1 | 0 | RP11-445F12.1 | 0 | ncRNA_intronic |
| rs61144937 | 17:35244495 | A | G | 0.47 | 21.51 | -0.09 | 0.02 | 3.50E-06 | RP11-445F12.1 | 0 | RP11-445F12.1 | 0 | ncRNA_intronic |
| rs11263809 | 17:35244752 | A | T | 0.47 | 21.52 | -0.09 | 0.02 | 3.50E-06 | RP11-445F12.1 | 0 | RP11-445F12.1 | 0 | ncRNA_intronic |
| rs72794140 | 16:83033038 | G | A | 0.92 | 21.19 | 0.16 | 0.04 | 4.20E-06 | CDH13 | 0 | CDH13 | 0 | intronic |
| rs34086136 | 19:19085090 | C | T | 0.53 | 21.18 | 0.09 | 0.02 | 4.20E-06 | RN7SL70P | 16023 | SUGP2 | 16607 | intergenic |
| rs4808170 | 19:19088342 | T | C | 0.53 | 21.04 | 0.09 | 0.02 | 4.50E-06 | RN7SL70P | 19275 | SUGP2 | 13355 | intergenic |
| rs6536772 | 4:164950164 | A | G | 0.69 | 20.77 | 0.09 | 0.02 | 5.20E-06 | MARCH1 | 0 | MARCH1 | 0 | intronic |
| rs12976783 | 19:19082898 | T | C | 0.53 | 20.72 | 0.09 | 0.02 | 5.30E-06 | RN7SL70P | 13831 | SUGP2 | 18799 | intergenic |
| rs56733468 | 7:70404411 | G | A | 0.82 | 20.60 | 0.11 | 0.03 | 5.70E-06 | RP11-575M4.1 | 101412 | WBSCR17 | 192744 | intergenic |
| rs72794139 | 16:83033003 | T | C | 0.91 | 20.57 | 0.16 | 0.03 | 5.70E-06 | CDH13 | 0 | CDH13 | 0 | intronic |
| rs6536775 | 4:164965629 | A | G | 0.69 | 20.48 | 0.09 | 0.02 | 6.00E-06 | MARCH1 | 0 | MARCH1 | 0 | intronic |
| rs7680457 | 4:164947015 | A | G | 0.32 | 20.33 | -0.09 | 0.02 | 6.50E-06 | MARCH1 | 0 | MARCH1 | 0 | intronic |
| rs72792145 | 16:83029723 | T | C | 0.91 | 20.21 | 0.15 | 0.03 | 6.90E-06 | CDH13 | 0 | CDH13 | 0 | intronic |

| | | | | | | | | | | | | | |
|---|---|---|---|---|---|---|---|---|---|---|---|---|---|
| rs72794156 | 16:83035565 | C | A | 0.92 | 20.19 | 0.15 | 0.03 | 7.00E-06 | CDH13 | 0 | CDH13 | 0 | intronic |
| rs10857366 | 4:164955338 | G | A | 0.68 | 20.13 | 0.09 | 0.02 | 7.30E-06 | MARCH1 | 0 | MARCH1 | 0 | intronic |
| rs1874581 | 17:35245000 | G | A | 0.46 | 20.09 | -0.09 | 0.02 | 7.40E-06 | RP11-445F12.1 | 0 | RP11-445F12.1 | 0 | ncRNA_intronic |
| rs35660429 | 7:70404645 | C | T | 0.34 | 20.05 | -0.09 | 0.02 | 7.50E-06 | RP11-575M4.1 | 101646 | WBSCR17 | 192510 | intergenic |
| rs11945181 | 4:164946487 | T | G | 0.32 | 19.99 | -0.09 | 0.02 | 7.80E-06 | MARCH1 | 0 | MARCH1 | 0 | intronic |
| rs8106237 | 19:19091076 | G | A | 0.54 | 19.89 | 0.09 | 0.02 | 8.20E-06 | RN7SL70P | 22009 | SUGP2 | 10621 | intergenic |
| rs35264282 | 10:5058490 | A | G | 0.71 | 19.87 | -0.10 | 0.02 | 8.30E-06 | AKR1C2 | 0 | AKR1C2 | 0 | intronic |
| rs36026148 | 10:5058493 | C | T | 0.71 | 19.86 | -0.10 | 0.02 | 8.30E-06 | AKR1C2 | 0 | AKR1C2 | 0 | intronic |
| rs13114716 | 4:164969077 | G | T | 0.68 | 19.85 | 0.09 | 0.02 | 8.40E-06 | MARCH1 | 0 | MARCH1 | 0 | intronic |
| rs72794153 | 16:83035286 | C | G | 0.92 | 19.83 | 0.15 | 0.03 | 8.40E-06 | CDH13 | 0 | CDH13 | 0 | intronic |
| rs76416971 | 17:17975454 | A | G | 0.95 | 19.81 | -0.22 | 0.05 | 8.60E-06 | GID4 | 3736 | DRG2 | 15746 | intergenic |
| rs72794135 | 16:83032676 | G | A | 0.91 | 19.75 | 0.15 | 0.03 | 8.80E-06 | CDH13 | 0 | CDH13 | 0 | intronic |
| rs6828312 | 4:164950846 | G | A | 0.68 | 19.73 | 0.09 | 0.02 | 8.90E-06 | MARCH1 | 0 | MARCH1 | 0 | intronic |
| rs4560357 | 4:164962875 | C | A | 0.68 | 19.72 | 0.09 | 0.02 | 9.00E-06 | MARCH1 | 0 | MARCH1 | 0 | intronic |
| rs34805072 | 10:5058482 | T | C | 0.71 | 19.70 | -0.09 | 0.02 | 9.00E-06 | AKR1C2 | 0 | AKR1C2 | 0 | intronic |
| rs13149496 | 4:164951675 | C | T | 0.68 | 19.70 | 0.09 | 0.02 | 9.10E-06 | MARCH1 | 0 | MARCH1 | 0 | intronic |
| rs11732646 | 4:164961778 | G | A | 0.68 | 19.63 | 0.09 | 0.02 | 9.40E-06 | MARCH1 | 0 | MARCH1 | 0 | intronic |
| rs72792163 | 16:83031484 | G | A | 0.91 | 19.60 | 0.15 | 0.03 | 9.50E-06 | CDH13 | 0 | CDH13 | 0 | intronic |
| rs11935788 | 4:164956662 | C | A | 0.40 | 19.56 | -0.09 | 0.02 | 9.70E-06 | MARCH1 | 0 | MARCH1 | 0 | intronic |
| rs72792161 | 16:83031258 | A | G | 0.91 | 19.57 | 0.15 | 0.03 | 9.70E-06 | CDH13 | 0 | CDH13 | 0 | Intronic |
| rs72792159 | 16:83031178 | A | T | 0.91 | 19.55 | 0.15 | 0.03 | 9.80E-06 | CDH13 | 0 | CDH13 | 0 | intronic |
| rs72792164 | 16:83031509 | G | C | 0.91 | 19.55 | 0.15 | 0.03 | 9.80E-06 | CDH13 | 0 | CDH13 | 0 | intronic |
| rs117212563 | 8:82519733 | C | T | 0.95 | 19.53 | 0.20 | 0.04 | 9.90E-06 | IMPA1P | 0 | IMPA1P | 0 | ncRNA_intronic |
| rs72792158 | 16:83031136 | G | A | 0.91 | 19.54 | 0.15 | 0.03 | 9.90E-06 | CDH13 | 0 | CDH13 | 0 | intronic |
